# Genome-wide analysis of heat stress-stimulated transposon mobility in the human fungal pathogen *Cryptococcus deneoformans*

**DOI:** 10.1101/2022.06.10.495668

**Authors:** Asiya Gusa, Vikas Yadav, Cullen Roth, Jonathan D. Williams, Eva Mei Shouse, Paul Magwene, Joseph Heitman, Sue Jinks-Robertson

**Author notes:** Sue Jinks-Robertson, **Email**. **Author Contributions**: A.G., J.D.W., V.Y., C.R, P.M., J.H. and S.J.R. designed research; A.G., J.D.W., V.Y., C.R, and E.S. performed research; A.G., V.Y., C.R, E.S., P.M., J.H., and S.J.R. analyzed the data; A.G. and S.J.R. wrote the paper. **Competing Interest Statement**: None.

## Abstract

We recently reported transposon mutagenesis as a significant driver of spontaneous mutations in the human fungal pathogen *Cryptococcus deneoformans* during murine infection. Mutations caused by transposable element (TE) insertion into reporter genes were dramatically elevated at high temperature (37° versus 30°) in vitro, suggesting that heat stress stimulates TE mobility in the Cryptococcus genome. To explore the genome-wide impact of TE mobilization, we generated transposon accumulation lines by in vitro passage of *C. deneoformans* strain XL280α for multiple generations at both 30° and at the host-relevant temperature of 37°. Utilizing whole-genome sequencing, we identified native TE copies and mapped multiple *de novo* TE insertions in these lines. Movements of the T1 DNA transposon occurred at both temperatures with a strong bias for insertion between gene-coding regions. By contrast, the Tcn12 retrotransposon integrated primarily within genes and movement occurred exclusively at 37°. In addition, we observed a dramatic amplification in copy number of the Cnl1 (*C. neoformans* LINE-1) retrotransposon in sub-telomeric regions under heat-stress conditions. Comparing TE mutations to other sequence variations detected in passaged lines, the increase in genomic changes at elevated temperature was primarily due to mobilization of the retroelements Tcn12 and Cnl1. Finally, we found multiple TE movements (T1, Tcn12 and Cnl1) in the genomes of single *C. deneoformans* isolates recovered from infected mice, providing evidence that mobile elements are likely to facilitate microevolution and rapid adaptation during infection.

**Significance Statement:** Rising global temperatures and climate change are predicted to increase fungal diseases in plants and mammals. However, the impact of heat stress on genetic changes in environmental fungi is largely unexplored. Environmental stressors can stimulate the movement of mobile DNA elements (transposons) within the genome to alter the genetic landscape. This report provides a genome-wide assessment of heat stress-induced transposon mobilization in the human fungal pathogen Cryptococcus. Transposon copies accumulated in genomes more rapidly following growth at the higher, host-relevant temperature. Additionally, movements of multiple elements were detected in the genomes of cryptococci recovered from infected mice. These findings suggest that heat stress-stimulated transposon mobility contributes to rapid adaptive changes in fungi both in the environment and during infection.

## Introduction

Rapid adaptations to stress conditions are vital for the survival and virulence of human fungal pathogens including those within the Cryptococcus species complex (1). Cryptococci are widely distributed in the environment and can be found as budding yeast or spores in soil, trees, and pigeon guano (2). Though cryptococci are naturally adapted for growth in the environment, the inhalation of spores can lead to human infection, particularly in individuals who are immunocompromised or immunosuppressed. Cryptococcus spp. can cause a lethal meningoencephalitis if the infection spreads systemically from the lungs and breaches the blood-brain barrier (3). Annually, cryptococcal meningitis afflicts nearly a quarter of a million people, causing 112,000 fatalities worldwide (4). Cryptococcosis is particularly deadly for HIV-infected populations, where it is estimated to be responsible for up to 19% of AIDS-related deaths (4). Serotypes A-D of Cryptococcus were originally defined by their capsular components and are now commonly recognized as distinct species by phylogeny (1). Infections by *Cryptococcus neoformans* (serotype A) are the most prevalent and lethal, primarily impacting HIV-infected populations in sub-Saharan Africa (5). Infections by the closely related *C. deneoformans* (serotype D) are less common and more typically associated with skin infections in European populations (6). Alarmingly, *C. gattii* species (serotypes B and C) can infect healthy, immunocompetent individuals, in addition to those with weakened immune systems (7). These infections are endemic in tropical and subtropical regions of the world, and cases also occur in British Columbia, Canada and the US Pacific Northwest (8, 9).

The transition from the environment to a warm-blooded host requires rapid adaptation to a variety of stresses including high temperature. For Cryptococcus spp., and other pathogenic fungi, the ability to tolerate heat stress is critical for virulence; those fungi unable to withstand high temperatures are unable to cause disease (10). Thermal stress induces global changes in the transcriptome and proteome that enable Cryptococcus to survive in the host. The cellular response to heat stress includes activation of target genes through the calcineurin-Crz1 pathway (11), protein stabilization by Hsp90 and other molecular chaperones (12) and the rapid decay of ribosomal protein mRNA as part of a generalized stress response (13).

In addition to programmed cellular responses to stress, spontaneous genomic changes that arise in the context of human fungal infections have been described (14). Adaptive genomic changes that confer drug resistance or enhance key virulence factors include changes in ploidy or gene copy number, as well as single nucleotide polymorphisms (SNPs) and minor insertions/deletions (INDELs) that alter gene expression or function (15–17). In addition, during lung infection, cryptococci can form giant polyploid cells, known as titan cells, that are resistant to phagocytosis and antifungal drug treatment (18, 19).

Spontaneous changes in eukaryotic genomes can also be mediated by transposable elements (TEs). TEs are “selfish” DNA elements that can insert within or between gene-coding regions to disrupt gene function or alter gene expression. In addition, TE sequences can serve as sites for recombination, facilitating genomic duplications, deletions and chromosomal rearrangements. There are two primary classes of TEs: 1) the DNA transposons, whose “cut and paste” movements are mediated by the element’s transposase, and 2) the retrotransposons, whose “copy and paste” movements require reverse transcription of an RNA intermediate mediated by enzymes encoded within the element, followed by integration of the element at a second site in the genome (20). Although TE movement is typically suppressed in the host genome, some elements can mobilize under conditions of stress. Stress-stimulated TE mobility in eukaryotes has been studied most extensively in plants and in Drosophila, where a number of abiotic stresses (UV radiation, wounding, temperature, nitrate starvation) and biotic stresses (viral or fungal infection) that increase TE mobility have been identified (21–24). Studies in budding and fission yeast have provided further evidence that environmental stimuli such as cold temperature or low oxygen conditions can trigger TE movement in the genome (25, 26).

Transposons make up ~5 to 6% of the Cryptococcus genome and the majority are retrotransposon sequences located in the centromeres of each chromosome (27, 28). A comparative study of the centromeres of three pathogenic species (*C. neoformans*, *C. deneoformans* and *C. deuterogattii*) found them to be devoid of open reading frames (ORFs), poorly transcribed, and rich in the sequences of retroelements Tcn1-Tcn6 (29). While the centromere-associated retroelements in *C. deuterogattii* are truncated and therefore non-mobile, *C. neoformans* and *C. deneoformans* centromeres additionally contain full-length and potentially active transposons whose movements are likely suppressed by heterochromatin and RNA interference (RNAi) silencing mechanisms.

Active transposition during mitotic growth has been reported in several *C. deneoformans* strains and most recently in *C. neoformans* (30–33). Two clinical strains of *C. neoformans* from South Africa were identified as hypermutators due to loss of RNAi function, which enabled mobilization of the Cnl1 (*C. neoformans* LINE-1) retrotransposon (33). In addition to a massive accumulation of Cnl1 copies in the sub-telomeric regions of chromosomes, Cnl1 inserted into the *FRR1* reporter gene (encoding FKBP1), resulting in rapamycin+FK506 drug resistance. Other reports have documented TE movements in *C. neoformans* during mating when the RNAi silencing machinery is inactivated or during mitotic growth when key components of the RNAi pathway are disabled (34, 35).

Recently, we reported TE-mediated drug resistance in clinical and environmental strains of *C. deneoformans* (31). Spontaneous mutants resistant to 5-fluoroorotic acid (5FOA), the antifungal drug 5-fluorocytosine, or the combination of antifungal drugs rapamycin+FK506, contained TE insertions in the *URA3/URA5*, *UXS1*, or *FRR1* genes, respectively. These TE insertions, which included a DNA transposon (T1) and a retrotransposon (Tcn12), were the primary cause of drug resistance at the higher, host-relevant temperature of 37° in vitro. Utilizing the *URA3/URA5* reporter genes, we found the temperature-dependent mobility of T1 and Tcn12 to be largely independent of RNAi control (31). Furthermore, T1 and Tcn12 insertions within *URA3/URA5* were found in the majority of 5FOA-resistant mutants recovered from mice following infection by the *C. deneoformans* XL280α strain.

The heat stress-induced mobilization of TEs in *C. deneoformans* suggests that this may be an adaptive strategy for enhanced survival and pathogenesis (31). However, the reporter genes previously utilized to identify TE insertions represent less than 5 kb of a 19 Mb genome. To characterize genome-wide TE movement, and to identify additional TEs that mobilize during conditions of thermal stress, we generated XL280α transposon accumulation (TA) lines at both 30° and 37°. Because the movements of TEs are often masked in whole-genome sequencing studies (36, 37), we applied a three-tiered approach to identify genome-wide *de novo* insertions, including Southern analysis, Illumina short-read sequencing with a custom TE mapping pipeline, and Nanopore long-read sequencing. Utilizing these approaches, we confirmed T1 and Tcn12 movement in passaged lines, but only Tcn12 mobility exhibited strong temperature dependence. T1 inserted predominantly between genes whereas Tcn12 inserted primarily within genes. We also observed significant mobilization of the Cnl1 retrotransposon at elevated temperature, with *de novo* copies of Cnl1 localized to sub-telomeric regions. In RNAi-deficient TA lines, genome-wide movements of T1 and Cnl1 were elevated at both 30° and 37°; Tcn12 movements, however, were independent of RNAi control. During murine infection all three elements mobilized in XL280α, with evidence of multiple TE changes in the genomes of recovered isolates. Finally, we compared the number of SNPs, INDELs and TE copy number changes in Illumina-sequenced lines and found that TE copy number changes drive the temperature-dependent increase in genomic changes at elevated temperature. Taken together, these data suggest that endogenous mobile elements have the potential to rapidly change the genomic landscape in response to heat stress within the environment and during host infection.

## Results

### Native copies of mobile T1 and Tcn12 elements outside of centromeres do not contain CpG methylation

The T1 DNA transposon and Tcn12 retrotransposon were previously identified as heat-mobile elements using the *URA3/URA5* reporters (31). To determine the endogenous chromosomal locations and copy numbers of these elements in *C. deneoformans* strain XL280α, the genome was assembled using Nanopore long-read and Illumina short-read sequencing (38). There were five 2.7 kb copies of T1 in the genome (two copies on chromosome 4, and one each on chromosomes 1, 5 and 6), with each copy positioned between predicted coding regions (Fig. 1 and *SI Appendix*, Fig. S1*A*). These elements share more than 98% sequence homology with the T1 sequence originally identified in *C. deneoformans* MCC48 (GenBank accession no. AY145841). Each T1 copy contained 11-bp terminal inverted repeats (TIRs), the sequence and length of which is transposon specific. TIRs serve as unique binding sites for the element’s transposase, which both excises and integrates the transposon into a new genomic location (39). The 2.7 kb T1 elements may be non-autonomous as they do not encode an ORF containing the DDE/DDD catalytic domain characteristic of transposases (40). There is, however, a putative ORF (CNE03210) annotated in the *C. deneoformans* JEC21 reference genome that is identical to that within the T1 element on chromosome 5 (*SI Appendix*, Fig. S1*B*). This ORF is predicted to encode a small 258 amino acid protein of unknown function, with a single coil domain in its N-terminus (InterPro). The corresponding transcript spans ~50% of the T1 sequence and was detected in RNA-seq data from XL280α cultures (41), but whether the putative protein is involved in T1 transposition is unknown.

**Figure 1.**
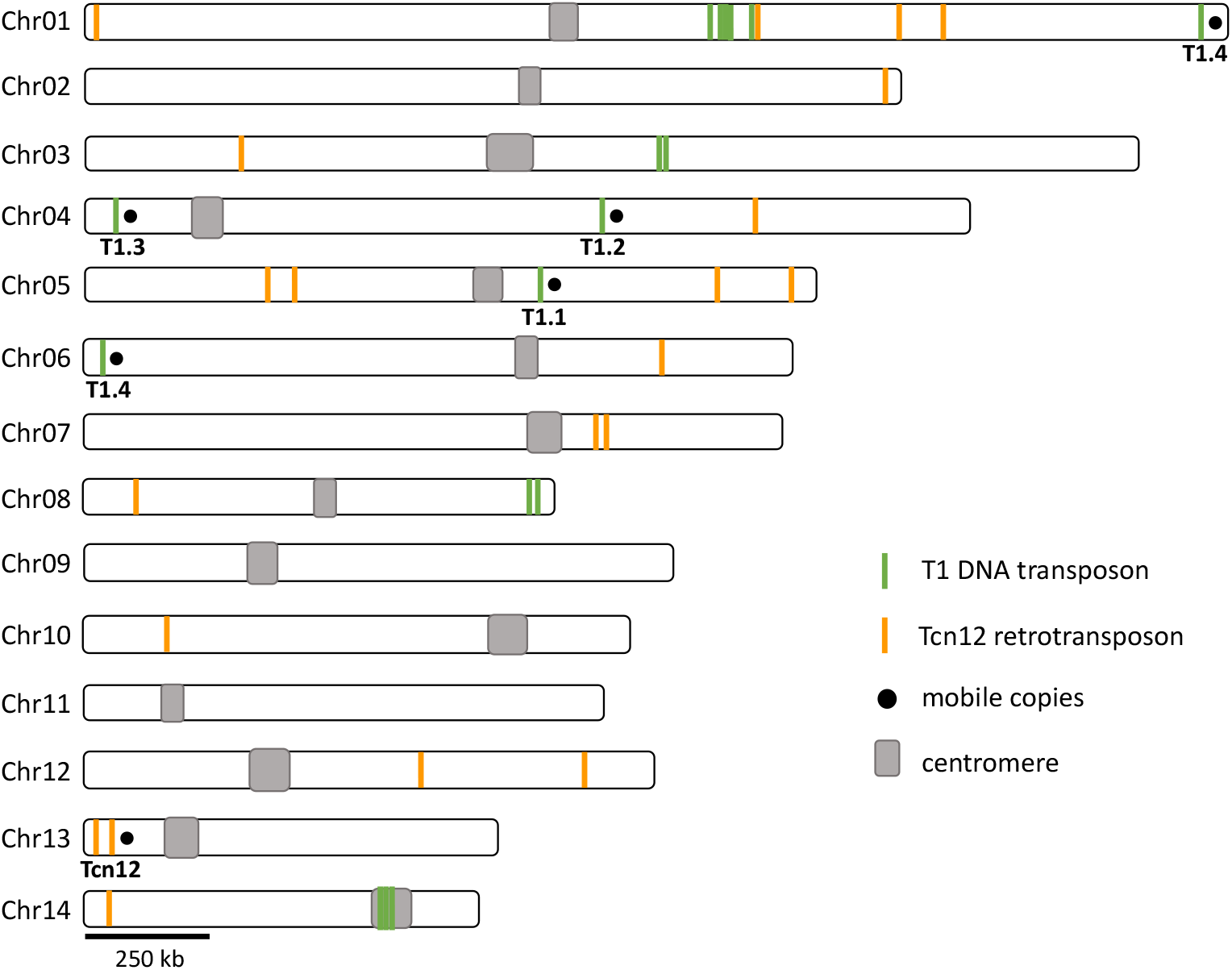
Location and copy number of full-length and truncated T1 and Tcn12 sequences in the *C. deneoformans* XL280α genome.

Four of the five T1 copies (T1.1 through T1.4) can be differentiated by minor SNPs or INDELs (*SI Appendix*, Fig. S1*B*). Each unique T1 sequence was found in the *URA3* or *URA5* gene of 5FOA drug-resistant mutants following growth in culture at 37°, demonstrating the mobility of each (*SI Appendix*, Fig. S1*C*)(31). In addition to the “full-length” 2.7 kb T1 elements, twelve truncated copies 0.15 kb to 1.7 kb in length with >90% sequence homology were identified throughout the genome (Fig. 1). Because no truncated copies of T1 were identified in the reporter genes that have captured novel TE insertions (*URA3*/5, *FRR1* or *UXS1*) in XL280α, they have presumably lost the ability to mobilize; each truncated T1 sequence lacked one or both of the TIR sequences required for recognition by the transposase.

A single, 5.2 kb full-length copy of Tcn12 was found on chromosome 13 in XL280α (Fig. 1). This copy encodes the reverse transcriptase and integrase required for movement, and has 152-bp long terminal repeats (LTRs) that are a characteristic feature of LTR retrotransposons (*SI Appendix*, Fig. S2*A*). In addition to the full-length copy, seven truncated Tcn12 sequences and thirteen copies of the Tcn12 solo LTR were scattered throughout the genome (Fig. 1). The solo LTR sequences presumably reflect homologous recombination between the LTR direct repeats (42). Tcn12 in XL280α shares 99% sequence identity to the Tcn12 sequence originally identified in the *C. neoformans* H99 reference genome (GenBank accession no. JQ277268).

In XL280α, as in other *C. deneoformans* and *C. neoformans* genomes, the majority of retrotransposon sequences (Tcn1-6) are located in the centromeric regions of each chromosome (29), with the notable exception of the Cnl1 non-LTR retrotransposon, which is located in the sub-telomeric regions (33, 43). Transposons in the centromeres are thought to be inactive due to silencing of repetitive elements by RNAi, heterochromatin, and DNA methylation (29, 34, 44). Notably, Tcn12 and all mobile copies of T1 within the XL280α genome were located outside of centromere regions.

Cryptococcus has no *de novo* DNA methyltransferase but does encode a single DNA methyltransferase (Dnmt5) that maintains 5-methylcytosine (5mC) DNA modification during mitotic growth (45). As a result, it is difficult to dynamically regulate the mobilization of transposons through this modification. However, it is possible that the temperature somehow alters the methylation landscape to allow transposition. From Nanopore sequence reads, we thus examined the XL280α genome for 5mC-methylation of CpG residues (*SI Appendix*, Fig. S3*A*). 5mC-methylation was most prevalent at predicted centromere sequences and at the sub-telomeres of some chromosomes, coinciding with the presence of repetitive elements, as previously shown for other Cryptococcus species (29, 45, 46). Importantly, we found that genomic DNA encoding the mobile T1 and Tcn12 elements was not methylated at CpG residues (*SI Appendix*, Fig. S3*B*). Therefore, the mobility of these elements is not likely to be controlled through a loss of methylation maintenance.

### Genomes of TA lines passaged at higher temperature accumulate more TE copies

To analyze novel TE integration sites in the presence or absence of heat stress, we generated TA lines by single colony passage of XL280α on non-selective medium 40 times (~800 generations total for each line). Forty-two independent TA lines (20 lines passaged at 30° and 22 lines passaged at 37°) were evaluated by Southern analysis and/or whole-genome sequencing. In Southern analysis using a probe specific for T1 (*SI Appendix*, Fig. S2*A*), eight fragments were detected in the “parental” unpassaged genome (Fig. 2*A* and *SI Appendix*, S2*B*). These fragments correspond to full-length as well as truncated T1 copies. Southern analysis with an internal probe specific for Tcn12 detected a single band in the parent, corresponding to the single full-length genomic copy (*SI Appendix*, Fig. S2*C*). T1 or Tcn12 movement was detected by the presence or absence of fragments compared to the unpassaged strain (*SI Appendix*, Fig. S2*B* and S2*C*). For example, Southern analysis of TA line 30-02, passaged at 30° (Fig. 2*A*), showed three new bands and the loss of one band compared to the unpassaged parent, indicating three novel T1 insertions and the loss of one copy from its native location.

**Figure 2.**
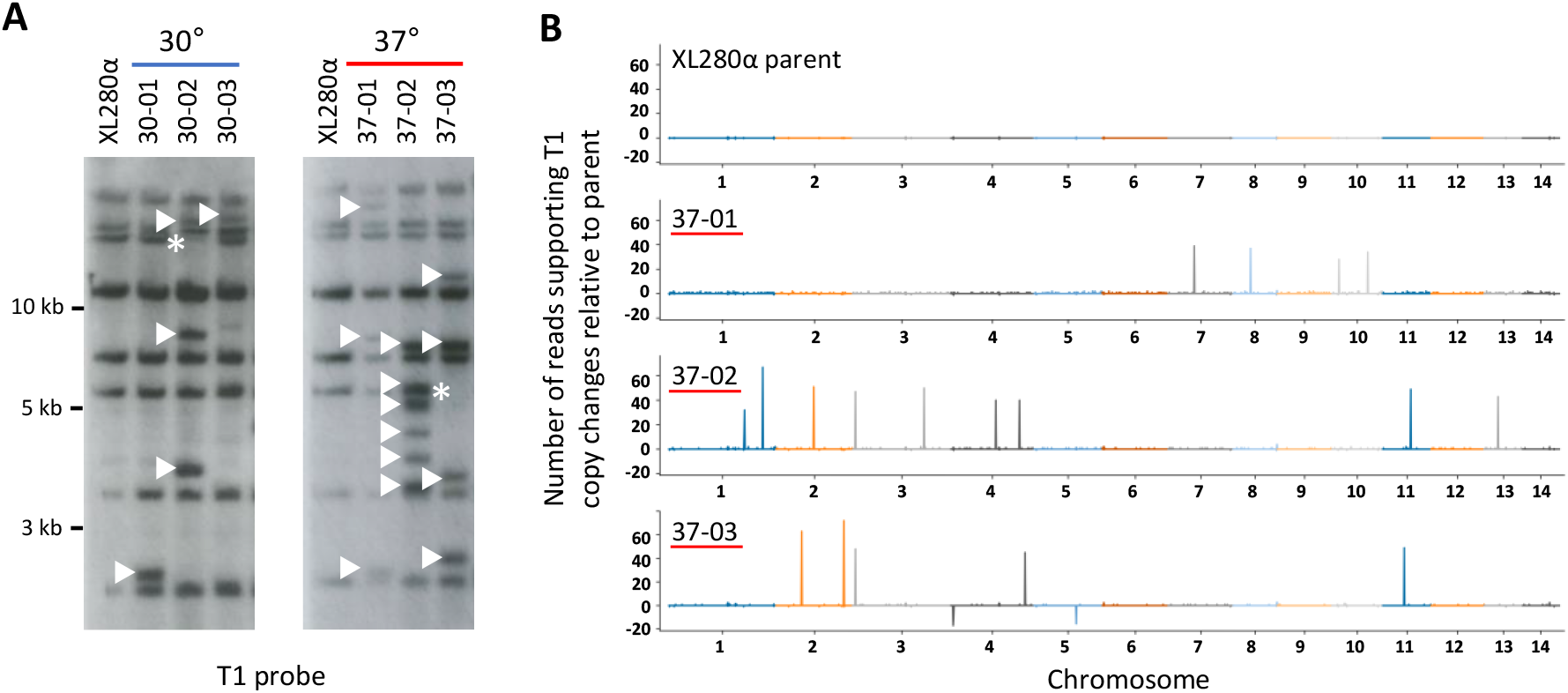
Mapping of TE changes in XL280α TA lines passaged ~800 generations. *(A)* Southern analysis of BspEI-digested genomic DNA probed for T1 (see *SI Appendix*, Fig. S2*A*) in TA lines passaged at 30° (30-01, 30-02, and 30-03) and 37° (37-01, 37-02, and 37-03). Arrowheads indicate *de novo* T1 insertions and asterisks indicate the loss of T1 copies relative to the parent genome. *(B)* T1 mapping using Illumina read-pair analysis. Read-depths for T1 in the passaged lines were normalized against the XL280α parent genome (top) to indicate insertion of new T1 copies (peaks) or the loss of T1 copies (troughs) in representative TA lines. The variation in height of each peak or trough corresponds to the number of supporting reads from the read-pair analysis.

To map the locations of *de novo* T1 and Tcn12 insertions in TA lines and to quantify copy number changes, Illumina pair-end sequencing was performed on eight lines passaged at 30° and ten lines passaged at 37°. Reads were analyzed to identify discordant pairs that mapped to both the genome and to the TE of interest (see methods). Briefly, all sequenced reads were mapped to the relevant TE, and the paired mates of these reads, which did not map to the TE, were used to anchor the position of the TE along a chromosome. Across the genome, windows of 10 kb were then analyzed for the enrichment of anchor reads. This analysis was also performed on the sequencing data of the non-passaged XL280α strain to generate the genome-wide background TE signal of anchoring reads. No evidence of aneuploidy in TA lines was found upon examining the genome-wide sequencing depth of pair-end reads (*SI Appendix*, Fig. S4).

Fig. 2*B* displays the new copies of T1 detected with the above strategy for three TA lines passaged at 37°. Not surprisingly, discordant read-pair analysis detected *de novo* T1 insertions or T1 copy loss at native sites that were not detected by Southern analysis. For example, four *de novo* T1 copies and the loss of one native T1 copy were detected by Southern analysis of TA line 37-03 (Fig. 2*A*), whereas discordant read-pair analysis of Illumina-sequenced reads detected five *de novo* T1 copies and the loss of two native T1 copies. All new T1 or Tcn12 copies as well as T1 losses predicted by read-pair analysis were confirmed by PCR, and Sanger sequencing was utilized to determine the exact locations of TE insertions (*SI Appendix*, Table S1). Additionally, we performed Nanopore long-read sequencing and genome assembly for seven of the TA lines, five of which had previously been sequenced using Illumina (30-01, 30-02, 37-01, 37-02, and 37-03). For the latter genomes, Nanopore sequencing detected two additional TE copies: a Tcn12 insertion within Cnl1 on chromosome 8 (TA line 37-01) and a T1 insertion adjacent to the T1.4 native copy on chromosome 6 (TA line 37-03). Compiling the sequencing data from 20 passaged genomes (13 sequenced by Illumina only, two sequenced by Nanopore only, and five sequenced using both platforms), we identified the locations of 46 novel T1 insertions and 19 novel Tcn12 insertions (Fig. 3*A* and 3*B* respectively, *SI Appendix*, Table S1). Although the locations of *de novo* T1 and Tcn12 copies were distributed fairly randomly across the genome, there were several *de novo* T1 copies in close proximity to the T1.4 native copy on chromosome 6 (Fig. 3*A*).

**Figure 3.**
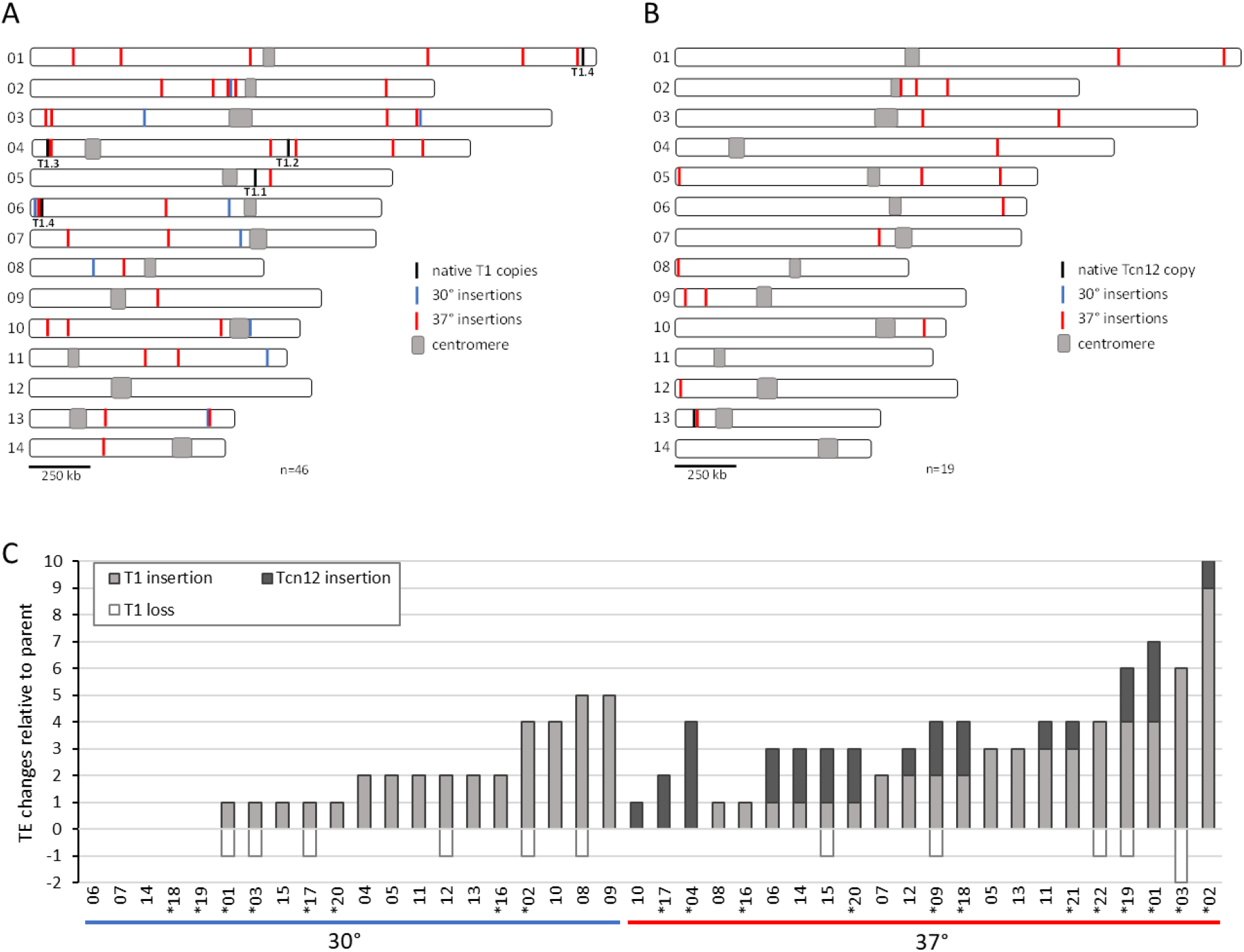
T1 and Tcn12 movement in TA lines passaged at 30° and 37° compared to the XL280α parent genome. *(A-B)*: Composite distribution of *de novo* T1 insertions (n=46) and Tcn12 insertions (n=19) mapped on chromosomes 1-14 by whole-genome sequencing in eight lines passaged at 30° and 12 lines passaged at 37°. *(C)* Total TE changes detected in TA lines for 20 genomes passaged at 30° (left) and 22 genomes passaged at 37° (right), evaluated using Southern analysis or whole-genome sequencing; sequenced genomes are indicated by asterisks.

Data from Southern and whole-genome sequencing analyses were combined to estimate the accumulation of T1 and Tcn12 insertions per TA line (Fig. 3*C*); for TA lines with TEs identified by more than one method, Nanopore or Illumina sequencing data were used (in that order). For the 20 TA lines passaged at 30° (~16,000 total generations), there were 35 novel TEs identified and only T1 movement was detected at this temperature (0.0022 TEs per generation). For the 22 TA lines passaged at 37° (~17,600 generations), the number of new copies of T1 (53) and Tcn12 (28) combined was 0.0046 TEs per generation. The 2.1-fold increase in the proportion of new TE copies at 37° compared to 30° is significant (*p*<0.0002) and this difference was primarily due to Tcn12 movement, which was observed only at higher temperature. Comparing the number of *de novo* insertions in lines passaged at 37° vs 30°, the difference is highly significant for Tcn12 (Mann-Whitney test, *p*<0.0001) but not for T1 (*p*=0.32).

Interestingly, native copies of T1 were preserved in most passaged lines with *de novo* T1 insertions; the loss of native copies was evident in only 11 of 34 genomes with novel T1 insertions (Fig. 3*C*). This was somewhat unexpected due to the “cut and paste” mechanism of DNA element movement. However, models for replicative transposition of DNA transposons have been proposed (47).

### T1 inserts predominantly between genes and can affect gene expression

In examining the locations of *de novo* T1 insertions in passaged TA lines, we found no evidence of conserved residues or a common sequence motif at integration sites, consistent with the random distribution of T1 insertions mapped previously in the *URA3/5* loci (31). However, we noted that the majority of insertions (41 of 46; 89%) were intergenic (*SI Appendix*, Table S1). Since ~85% of the Cryptococcus genome is predicted to be coding, and only 15% non-coding (28), this suggested a strong bias for T1 insertion between genes. Approximately 18% to 26% of genes in Cryptococcus are likely essential, based on estimates in yeast with a similar number of genes (*Saccharomyces cerevisiae* and *Candida albicans*) (48, 49). Therefore, excluding essential coding regions (~20%), we would expect 0.68 of insertions to occur within genes and 0.32 to occur between genes if there were no insertion bias. The observed number of intragenic and intergenic T1 insertions in the passaged lines (five and 41, respectively), indicates a strong bias for T1 insertion between genes (Chi-square goodness of fit, *p*<0.0001).

Interestingly, T1 insertions were found between genes in tandem or divergent orientation, 15 (37%) and 26 (63%), respectively, but none were between genes in convergent orientation (*SI Appendix*, Table S2, Fig. S5*A*). In the XL280α genome, the proportion of genes in tandem, divergent, and convergent orientation is 44%, 39%, and 17%, respectively (*SI Appendix*, Table S2, Fig. S5*B*); overlapping genes where there is no intergenic region for TE insertion were excluded. Comparing the observed frequency of T1 insertions between genes in each orientation with the expected frequency based on the proportion of genes in each orientation in the XL280α genome, we found a bias for insertion between divergent genes (Chi-square test of association, *p*=0.0030) and a bias against insertion between convergent genes (*p*=0.0081)(*SI Appendix*, Table S2). Additionally, since most intergenic T1 insertions (34 of 41) mapped within 200 bp of the nearest predicted transcription start site (*SI Appendix*, Fig. S5*C*), this suggests that T1 insertion may frequently affect gene expression from nearby promoters.

The potential impact of *de novo* T1 insertion on nearby gene expression was examined in TA lines 37-01 and 37-02 (Fig. 4). Quantitative RT-PCR was employed to measure the transcript levels of six divergently trancribed gene pairs located proximal to T1 insertion sites. Nine of the 12 genes analyzed had a significant increase in gene expression in the passaged line compared to the non-passaged parent (1.3-fold to 26-fold increase, blue vertical arrows in Fig. 4). To address whether changes in gene expression occurred in other genes not proximal to *de novo* T1 insertions, we assessed gene expression from sets of divergently transcribed control genes in each line. Compared to transcript levels in the unpassaged parent, there were also significant changes in gene expression for five of 12 control genes tested (1.7-fold to 6.0-fold increases, *SI Appendix*, Fig. S6). These data suggest that T1 insertion affects the expression of nearby genes, and that additional, more global effects on gene expression occurred in these lines.

**Figure 4.**
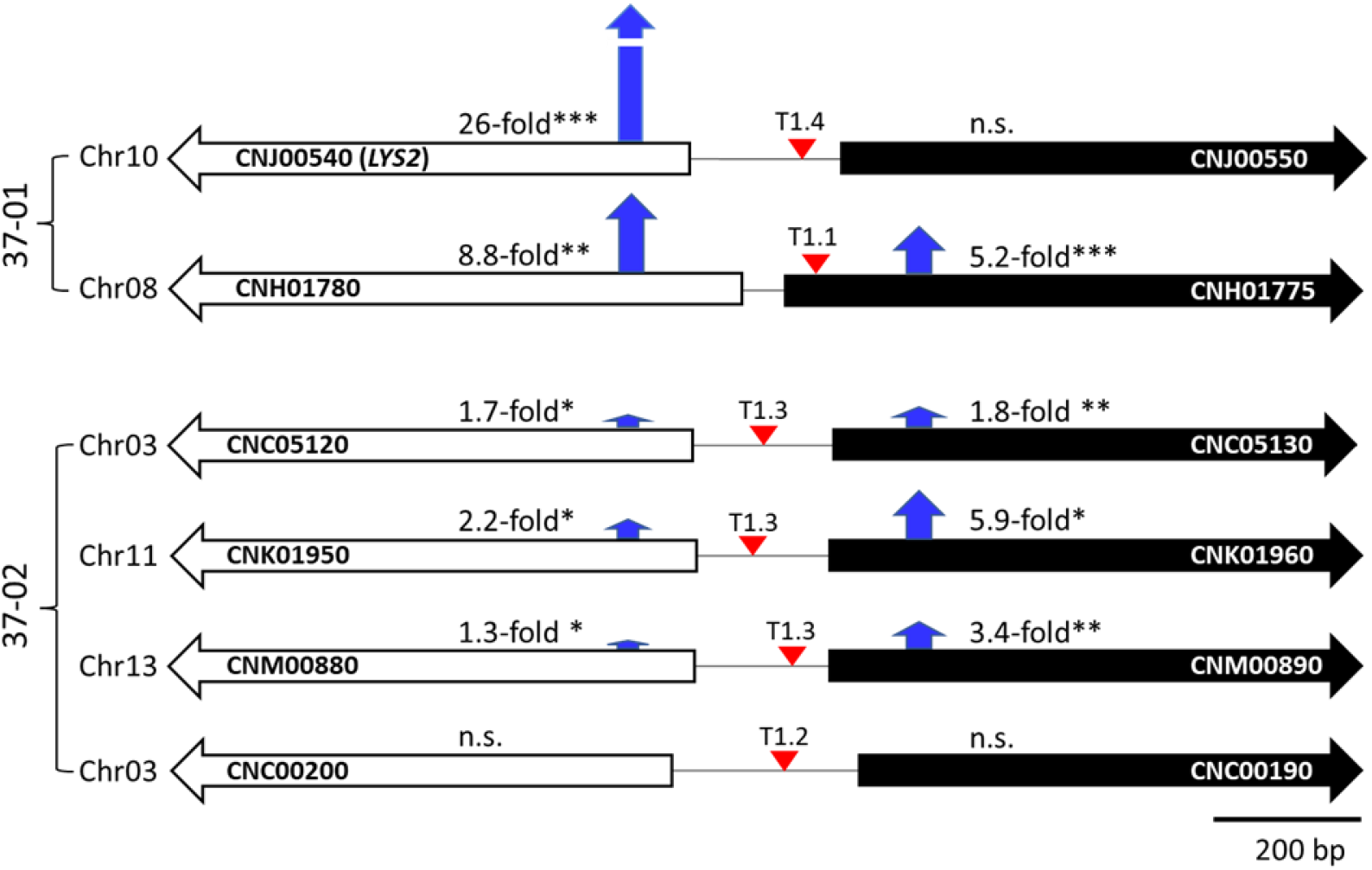
T1 insertions and changes in gene expression in 37°-passaged lines 37-01 and 37-02. Locations of *de novo* T1 insertions (red arrowheads) relative to predicted genes and fold changes in expression compared to the XL280α parent are shown. Black or white shading represents forward or reverse gene orientation, respectively, in the JEC21 reference genome (EnsemblFungi). Blue arrows indicate the relative fold-changes in transcript compared to the non-passaged XL280α strain measured by qRT-PCR. The fold change in expression was calculated using the comparative ΔΔC_T_ method. A Welch’s unpaired t test was performed for each pairwise comparison using the mean ΔC_T_ values for three biological replicates (n.s., not significant; * indicates 0.01<*p*≤0.05; and ** indicates 0.001<*p*≤0.01).

### Tcn12 inserts primarily in coding regions with conserved residues at integration sites

Of the 19 novel Tcn12 insertion sites mapped in sequenced TA lines passaged at 37°, 14 were found in coding regions (*SI Appendix*, Fig. S7). Other insertions mapped in non-coding regions between genes or near the centromere or telomere regions, with one insertion found within a Cnl1 element (*SI Appendix*, Table S1). Both the position and orientation of Tcn12 insertions appeared to be random. Although the majority of insertions were intragenic (74%), we did not find evidence for a Tcn12 insertion bias within genes based on the proportion (0.68) expected to insert randomly within non-essential coding regions in the genome (Chi-square goodness of fit, *p*= 0.76).

In previous analysis of *de novo* Tcn12 integration sites in the *URA3* and *URA5* reporters in XL280α, Tcn12 insertions occurred almost exclusively in *URA5*, indicating locus bias or specificity (31). Further, we identified a hotspot for Tcn12 insertions in the *URA5* ORF in which the element inserted multiple times in both orientations (*SI Appendix*, Fig. S8*A*). Where Tcn12 inserts, a 5-bp target-site duplication (TSD) occurs; the duplication size reflects the distance between integrase-generated staggered nicks that allow TE insertion, followed by the filling of small gaps and ligation (39). To assess whether there was sequence specificity for Tcn12 insertion genome-wide, we aligned sequences upstream and downstream of the TSD for seventeen *de novo* Tcn12 integration sites mapped in the TA lines (*SI Appendix*, Table S3); two integration sites did not have a discernable TSD and were therefore excluded. WebLogo (50) identified several conserved residues within a 15-bp sequence logo centered at the 5-bp TSD (*SI Appendix*, Fig. S8*B*). This sequence, ANGTN-TSD-NACNT, has palindromic features, with the most prominent residue conservation 2 to 3 bp from the TSD. Alignment of 21 unique Tcn12 integration sites in *URA5* from 5FOA-resistant cryptococci recovered from mice (31) yielded a similar, but weaker, palindromic sequence logo (*SI Appendix*, Fig. S8*C*). These data suggest a preferred binding motif for the Tcn12 integrase. For the *Ty1/copia*-like family of retrotransposons (which Tcn12 resembles), both DNA structure and sequence specificity appear to play a role in preferred target sites, as demonstrated for the *Tos17* and *Tto1* elements found in plant species (51, 52).

### Copies of the non-LTR retrotransposon Cnl1 increased in TA lines passaged at 37°

In analyzing the Nanopore data from the seven TA lines sequenced, we found a large increase in the number of full-length and truncated copies of the Cnl1 retrotransposon, primarily in lines passaged at 37° (Table 1, Fig. 5 and *SI Appendix*, Fig. S9). Cnl1 is the most abundant non-LTR retroelement in Cryptococcus and is typically located in sub-telomeric regions (43). In the parental XL280α genome, we identified nine full-length (3.4 kb) copies of Cnl1 (Fig. 5*A*); five of these copies on chromosomes 4, 5, 10 and 11 are not CpG-methylated and potentially mobile (*SI Appendix*, Fig. S3*C)*. Additionally, there were 67 truncated Cnl1 copies, located near the telomeres of most chromosomes (Fig. 5*A*). The greatest increase in Cnl1 copy number among the five sequenced lines passaged at 37° occurred in TA line 37-02, where the number of new, full-length copies of Cnl1 increased from nine to 49 copies (Table 1, Fig. 5*B*). New copies appeared on sub-telomeric arms of chromosomes where a native Cnl1 copy was present, and also on chromosomes where there were no existing copies (Fig. 5*B* and *SI Appendix*, Fig. S9). In addition to full-length copies, there was a considerable increase in truncated segments of Cnl1, most of which were truncated at the 5’ end. This pattern presumably reflects incomplete transposition events in which target-primed reverse transcription of the full-length element (3’ to 5’) at the integration site terminates prematurely (53). Notably, we did not find full-length Cnl1 copies in the genomes of the parental or passaged lines outside of telomeric regions, and many were inserted within pre-existing Cnl1 copies. Cnl1 was previously described to insert preferentially within itself in tandem orientation forming nested arrays that are usually found in proximity to proposed telomere repeat sequences in *C. neoformans* (33, 43). These nested arrays are consistent with the sub-telomeric localization of *de novo* Cnl1 arrays we observed in the passaged lines (Fig. 5*B* and *SI Appendix*, Fig. S9).

**Table 1.** Full-length TE copy numbers in XL280α TA lines sequenced by Nanopore^1^.

| TE copies | XL280α | 30-01 | 30-02 | 37-01 | 37-02 | 37-03 | 37-04 | 37-09 |
| --- | --- | --- | --- | --- | --- | --- | --- | --- |
| T1 | 5 | 5 | 8 | 9 | 14 | 9 | 5 | 6 |
| Tcn12 | 1 | 1 | 1 | 4 | 2 | 1 | 5 | 3 |
| Cnl1 | 9 | 9 | 12 | 37 | 49 | 22 | 33 | 41 |
| TOTAL | 15 | 15 | 21 | 50 | 65 | 32 | 43 | 50 |
| CHANGE | n/a | +0 | +6 | +35 | +50 | +17 | +28 | +35 |

<sup>1</sup>Lines passaged at 30° or 37° are indicated by 30-xx or 37-xx, respectively.

**Figure 5.**
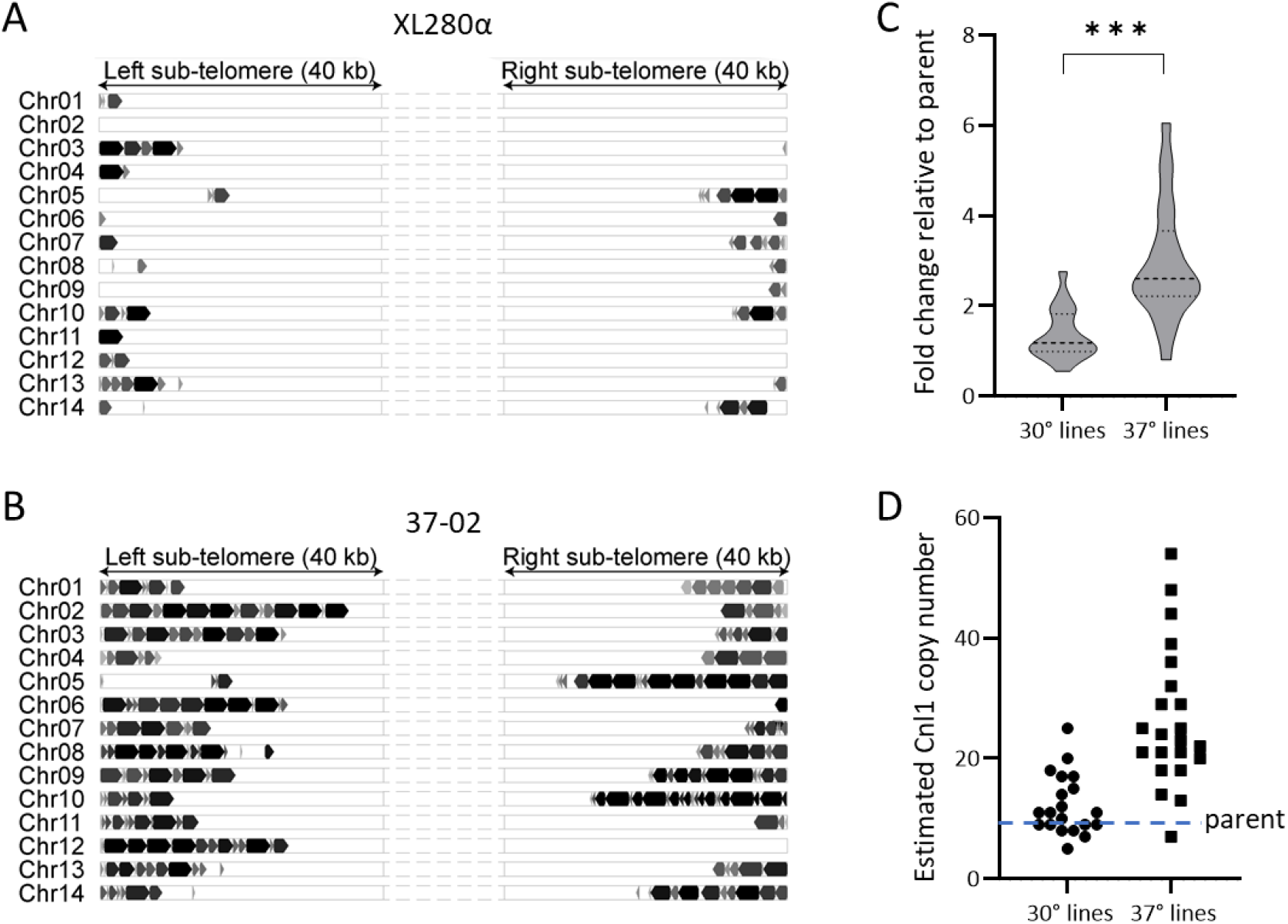
Cnl1 copy number changes in passaged lines. Full-length and truncated copies of Cnl1 in the parental XL280α genome *(A)* compared to TA line 37-02 *(B)*. Each panel *(A-B)* shows the sub-telomeric regions of each chromosomal arm. Cnl1 copies are represented by block arrows whose size approximates the length relative to the 3.4 kb full-length element. Full-length and truncated copies are indicated in black and gray, respectively. *(C)* Fold change in Cnl1 copy number for all TA lines passaged at 30° (n=20) and 37° (n=22) relative to the parent strain as determined by qPCR of the 5’-end of Cnl1 from genomic DNA. The middle dotted line of each violin plot represents the median fold change value while the top and bottom dotted lines represent the upper and lower quartile values, respectively. The fold change in copy number was calculated using the comparative ΔΔC_T_ method with *GPD1* as the internal control gene. The Mann-Whitney test was applied to compare the difference for each set of mean ΔC_T_ values from three biological replicates (*** indicates *p*<0.0001). *(D)* The estimated Cnl1 copy number for each TA line is plotted, determined by multiplying the fold change value by nine copies, which equals the number of full-length Cnl1 copies in the XL280α genome.

To confirm that the Cnl1 burden was higher in TA lines passaged at 37° compared to those passaged at 30°, we performed quantitative PCR of the 5’-end of Cnl1 in all TA lines to determine the fold change in copy number (Fig. 5*C*) and to estimate the number of full-length genomic copies (Fig. 5*D*). For TA lines passaged at 30°, the fold change in Cnl1 copies relative to the parent strain ranged from 0.56 to 2.8-fold, with a median value of 1.2 (11 copies). Fold changes in Cnl1 were more widely distributed among lines passaged at 37°, ranging from 0.82 to 6.1-fold, with a median value of 2.6 (23 copies). Comparing the C_T_ (cycle threshold) values for all lines passaged at 37° and 30°, we found a significant difference in the Cnl1 copy number for lines passaged at the higher temperature (Mann-Whitney, *p*<0.0001). Additionally, in Southern analysis probing for Cnl1 in the genomes of ten independent lines passaged at each temperature, we found that the hybridization signal present in the 37° lines was much stronger, particularly for higher molecular weight fragments (*SI Appendix*, Fig. S10). This is consistent with the pattern of nested Cnl1 arrays identified in the Nanopore data (Fig. 5, *SI Appendix*, Fig. S9).

### RNAi control of mobile elements is independent from temperature control

We previously found that temperature control of T1 and Tcn12 insertion into reporter genes was largely independent of RNAi silencing (31). To extend this analysis genome-wide, and to assess the impact of RNAi on the temperature-dependent mobility of Cnl1, we generated TA lines from an *rdp1*Δ derivative of XL280α following passage of single colonies on non-selective medium at 30° and 37°. By Southern analysis, multiple T1-specific fragments appeared or were lost in the absence of RNAi in TA lines passaged at 30° (*SI Appendix*, Fig. S11*A*); these movements were visibly more numerous compared to wildtype TA lines passaged at the same temperature (*SI Appendix*, Fig. S2*B*). At 37°, the T1 changes in most *rdp1*Δ lines were too numerous to count and obscured the nine-band hybridization pattern observed for the wildtype and *rdp1*Δ parent strains. This apparent increase in T1 mobility at 37° suggests that heat stress and the loss of RNAi have an additive or possibly synergistic effect on T1 mobility.

By contrast, loss of RNAi did not alter the mobility of Tcn12 in genomes passaged at 30° or 37°. Tcn12 movements were evident only in *rdp1*Δ lines passaged at 37° (*SI Appendix*, Fig. S11*B*), and the frequency of these movements in both the wildtype and *rdp1*Δ background was similar (~1.3 *de novo* copies per genome). For example, in wildtype TA lines passaged at 37°, 15 of 22 genomes contained new Tcn12 copies (Fig. 3*C*), and in *rdp1*Δ TA lines passaged at 37°, four of seven genomes assessed contained new Tcn12 copies (*SI Appendix*, Fig. S11*B*). There was no significant difference in the number of Tcn12 movements in the wildtype or *rdp1*Δ background (Fisher’s exact, *p*=0.46), indicating that RNAi does not suppress Tcn12 mobility at either temperature, consistent with previous findings (31).

Cnl1 mobility was shown to be controlled by RNAi in *C. deneoformans* JEC21 (30). In XL280α, Southern analysis of *rdp1*Δ TA lines passaged at 30° and 37° demonstrated a massive increase in Cnl1-specific fragments compared to wildtype passaged lines (*SI Appendix*, Fig. S11*C*), confirming that Cnl1 mobility is suppressed by RNAi in *C. deneoformans*. Although the bands indicating Cnl1 hybridization to *rdp1*Δ lines passaged at 37° appear somewhat darker than for *rdp1*Δ lines passaged at 30°, the difference was too subtle to gauge whether temperature stress at 37° stimulates Cnl1 movement beyond that observed at 30°. However, data from whole-genome sequencing and qPCR of wildtype TA lines passaged at 37° demonstrated that heat stress is sufficient to dramatically increase Cnl1 copy number when RNAi is functional (Table 1, Fig. 5).

### TEs drive the increase in genomic changes at elevated temperature

To compare the relative contribution of TE copy number changes to more commonly identified genomic changes (SNPs and INDELs), we analyzed the 18 Illumina-sequenced TA lines for sequence variations compared to the parental XL280α genome (see methods, *SI Appendix*). A total of 142 unique sequence variants were identified and characterized (*SI Appendix*, Table S4). Table 2 summarizes the number of genomic changes including SNPs, INDELs and *de novo* TE copies detected in each line. To determine the rate of events per generation, the total number of events for each category was divided by the number of generations in eight lines passaged at 30° (6400 generations) and ten lines passaged at 37° (8000 generations). The per generation rates of SNPs (0.0081) and INDELs (0.0022) in 30° lines were nearly identical to the rates in 37° lines (0.0078 and 0.0018, respectively) with no difference in the proportion of events that occurred at different temperatures (*p*=0.40, SNPs; *p*=0.28, INDELs). Extending this analysis per bp in the 19 Mb Cryptococcus genome, mutation rates for SNPs (4.1 to 4.2 × 10^−10^ per bp per generation) and for INDELs (0.92 to 1.2 × 10^−10^) in the passaged lines were comparable to mutation rates determined for other haploid yeast genomes (SNPs, 1.7 to 4.0 × 10^−10^; INDELs, 0.16 to 1.74 × 10^−10^)(54–56). By contrast, the proportion of new TE copies per generation was five-fold higher at 37° (0.031) compared to 30° (0.0061)(*p*<0.0002). This significant increase in TE copy number in lines passaged at 37° was due to the increased mobility of the retroelements Tcn12 and Cnl1 (Mann-Whitney test, p=0.0065 and p=0.0010, respectively). Additionally, the per generation rate of TE mutation at 37° was three times higher than the rate of SNPs and INDELs combined (0.031 compared to 0.0095). Taken together, these data indicate that TE mutations, rather than minor sequence variations, drive the increase in spontaneous genomic changes in *C. deneoformans* under conditions of heat stress.

**Table 2.** Comparison of sequence variations and *de novo* TE copies detected in XL280α TA lines passaged a total of ~6400 generations at 30° or ~8,000 generations at 37°C^1-3^.

| 30° lines | SNPs | INDELs | T1 | Tcn12 | Cnl1 | SNPs/<br>INDELs | Total<br>TEs |
| --- | --- | --- | --- | --- | --- | --- | --- |
| 30-01 | 5 | 0 | **1** | **0** | **0** | 5 | **1** |
| 30-02 | 6 | 2 | **4** | **0** | **3** | 8 | **7** |
| 30-03 | 2 | 1 | 1 | 0 | 2 | 3 | 3 |
| 30-16 | 7 | 3 | 2 | 0 | 0 | 10 | 2 |
| 30-17 | 6 | 1 | 1 | 0 | 2 | 7 | 3 |
| 30-18 | 6 | 3 | 0 | 0 | 16 | 9 | 16 |
| 30-19 | 8 | 2 | 0 | 0 | 6 | 10 | 6 |
| 30-20 | 12 | 2 | 1 | 0 | 0 | 14 | 1 |
| TOTAL | 52 | 14 | 10 | 0 | 29 | 66 | 39 |
| PER<br>GENERATION | 0.0081 | 0.0022 | 0.0016 | 0.000 | 0.0045 | 0.010 | 0.0061 |
| 37° lines | SNPs | INDELs | T1 | Tcn12 | Cnl1 | SNPs/<br>INDELs | Total<br>TEs |
| 37-01 | 8 | 2 | **4** | **3** | **28** | 10 | **35** |
| 37-02 | 5 | 3 | **9** | **1** | **40** | 8 | **50** |
| 37-03 | 7 | 0 | **6** | **0** | **13** | 7 | **19** |
| 37-16 | 17 | 1 | 1 | 0 | 14 | 18 | 15 |
| 37-17 | 4 | 1 | 0 | 2 | 16 | 5 | 18 |
| 37-18 | 4 | 1 | 2 | 2 | 9 | 5 | 13 |
| 37-19 | 7 | 4 | 4 | 2 | 20 | 11 | 26 |
| 37-20 | 5 | 1 | 1 | 2 | 15 | 6 | 18 |
| 37-21 | 2 | 1 | 3 | 1 | 30 | 3 | 34 |
| 37-22 | 3 | 0 | 4 | 0 | 12 | 3 | 16 |
| TOTAL | 62 | 14 | 34 | 13 | 197 | 76 | 244 |
| PER<br>GENERATION | 0.0078 | 0.0018 | 0.0043 | 0.0016 | 0.025 | 0.0095 | 0.031 |

<sup>1</sup>SNPs (single nucleotide polymorphisms), INDELs (1-13 bp insertions or deletions) and *de novo* T1 and Tcn12 copies were detected by Illumina whole-genome sequencing analyses.
<sup>2</sup>Cnl1 values represent estimated new full-length copies determined by qPCR.
<sup>3</sup>Bolded values indicate TE copies detected or confirmed by Nanopore sequencing.

### Multiple TEs mobilize in the genomes of cryptococci recovered from infected mice

To evaluate TE movements in *C. deneoformans* in vivo, we isolated genomic DNA from 5FOA-resistant colonies recovered from the organs of two mice ten days after intravenous infection with XL280α (31). Each of the nine mutants analyzed contained independent Tcn12 insertions in the *URA5* locus. In Southern analysis, the Tcn12 probe hybridized to an additional DNA fragment in each of the mutants recovered from mice (Fig. 6*A*), which presumably corresponds to the Tcn12 insertions in *URA5*. To examine whether genomic DNA from these recovered isolates had additional TE movements, we re-probed the same membrane for T1 and Cnl1 sequences (Fig. 6*B* and 6*C*, respectively). For three independent mutants recovered from the lungs, kidneys, and brain of infected mice (Fig. 6; lanes 2, 5 and 9, respectively), we also detected additional T1- and Cnl1-specific fragments. For example, in the single 5FOA-resistant mutant recovered from the brain of mouse 2 (lane 9), two movements of T1 (Fig. 6*B*) and at least three new Cnl1-specific fragments were detected (Fig. 6*C*).

**Figure 6.**
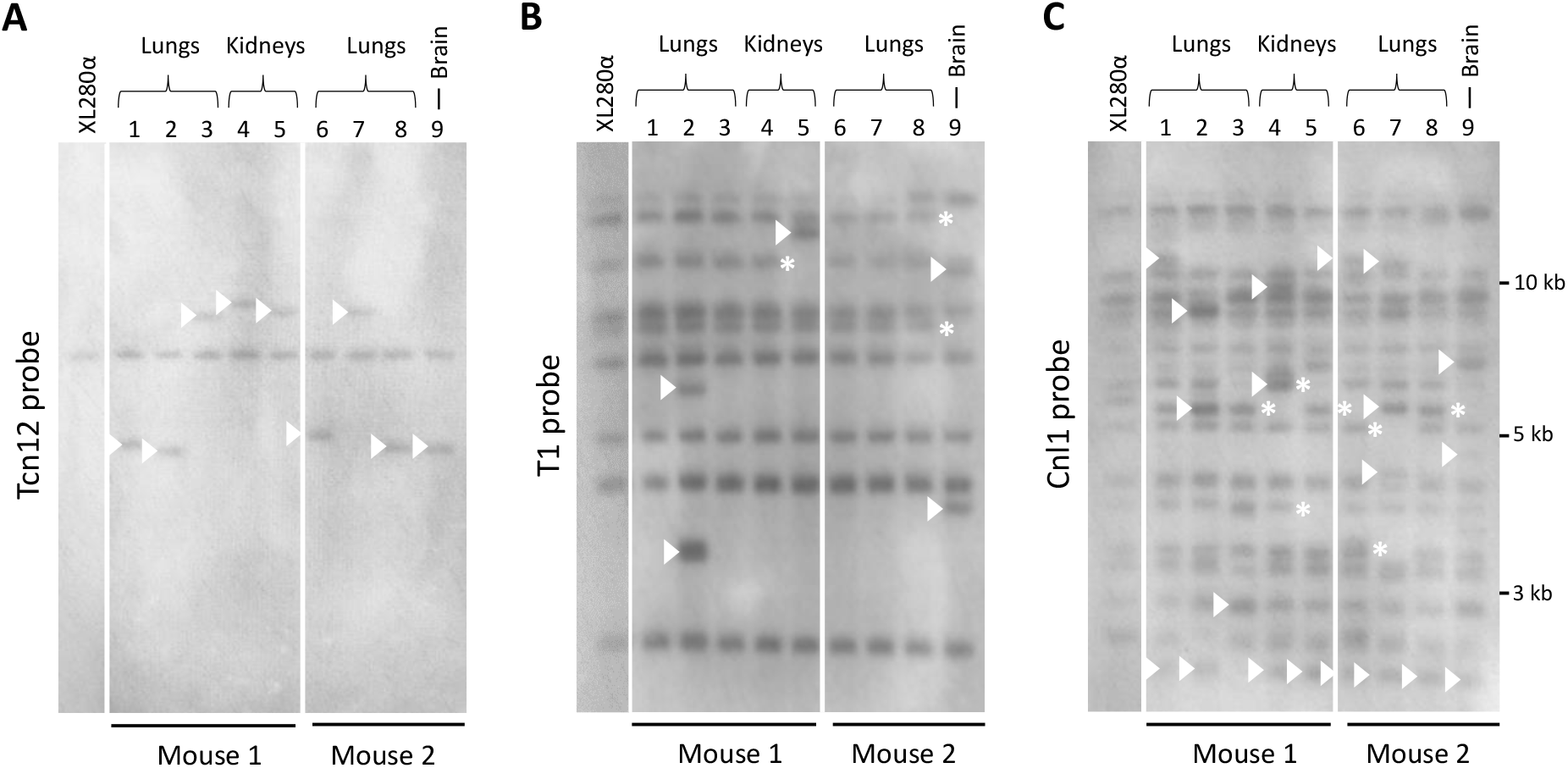
Southern analysis of TE fragments in 5FOA-resistant XL280α mutants recovered from the lungs, kidneys and brain of two mice ten days post-infection. Each mutant had an independent Tcn12 mutation in the *URA5* gene. Genomic DNA was digested with PvuII and probed for *(A)* Tcn12, *(B)* T1 or *(C)* Cnl1. The XL280α inoculum strain is shown for comparison. Arrows indicate new TE fragments and asterisks indicate TE fragment loss.

## Discussion

The survival of environmental fungi in mammalian hosts requires a rapid adaptation to elevated body temperature and the ability to replicate under conditions of sustained heat stress. Yet, little is known about the impact of heat stress on fungal evolution and whether stresses encountered in the environment or during infection stimulate genetic changes. Stress-stimulated genetic changes, such as those mediated by TE movements, may contribute to the evolution of pathogenic traits in fungi both in the environment and during infection.

In this study, we performed a genome-wide analysis of temperature-dependent TE movement in the human fungal pathogen *C. deneoformans* by evaluating independent TA lines passaged for 800 generations in vitro. *De novo* insertions of the T1 DNA transposon, the Tcn12 retrotransposon, or both, were detected in all 22 lines passaged at 37° and T1 movement was detected in 15 of 20 lines passaged at 30°. In addition, we identified movements of the Cnl1 non-LTR retrotransposon in sub-telomeric regions of chromosomes. Overall, we found a greater accumulation of TE copies in the genomes of TA lines passaged at the host-relevant temperature of 37° compared to those passaged at 30° (Tables 1 and 2). The increased TE copy number among 37° TA lines was due in part to mobilization of Tcn12, which occurred exclusively at elevated temperature in both wildtype and RNAi-deficient backgrounds. The greatest difference in copy number was detected for Cnl1; for 37° lines, the median copy number increased by 12 copies compared to a single copy increase in 30° lines.

One key finding from this study was that movements of all three elements (T1, Tcn12, and Cnl1) were detected in single genomes of Cryptococcus recovered from mouse organs (Fig. 6). This was surprising given the short incubation period of ten days in the host following intravenous infection. During this period, we estimated that a minimum of eight to ten generations of growth occurred (31). The actual number of in vivo generations is undoubtedly higher than this estimate, considering that some clearance of Cryptococcus by immune cells likely occurred. Although a direct comparison is not possible, only half of the TA lines (12 of 22) passaged at 37° in vitro had *de novo* T1 and Tcn12 movements in the same genome after 800 generations of non-selective growth on nutrient-rich medium (Fig. 3*C*). This suggests that TE mobilization within the host is more robust compared to growth in culture at 37°, and factors in addition to temperature stress (i.e. oxidative, nitrosative or pH stress) may increase TE mobility in vivo. Further, the rapid accumulation of TEs in some cryptococci recovered from mice suggests that there may be selection for TE-mediated adaptive mutations in a competitive growth environment. Alternatively, if stochastic bursts of transposition occur in single cells, we may have selected for mutants with multiple transposition events by evaluating only those isolates known to contain at least one TE mutation. The possibility that bursts of transposition occur in individual genomes is supported by the variation in the number of spontaneous transposition events detected among individual TA lines passaged at 30° and 37° (Fig. 3*C* and Table 1). Whether TE copy number changes occurred in single replication cycles or gradually over multiple generations has not been evaluated.

The temperature-dependent transposition of Tcn12 in the passaged genomes is consistent with previous results obtained with reporter genes (*URA3/5, FRR1* or *UXS1*), where Tcn12 insertions were detected at 37° and not at 30° (31). Unexpectedly, we found T1 movement on solid medium at 37° compared to 30°, was not significantly different (Fig. 3*C*, Mann-Whitney test, *p*=0.32), indicating that global mobilization of T1 is less temperature-dependent than previously inferred (31). In the prior study, which evaluated spontaneous 5FOA resistance in XL280α in liquid culture, the rate of T1 insertions in *URA3/5* at 37° was ~200-fold higher than at 30°. The reason for the T1-specific discrepancy is not clear, but could reflect a difference due to growth conditions (liquid versus solid medium) or an anomaly of the reporter genes used previously.

A second key finding from this study highlights the limitations of using reporter genes to detect TE movements in the genome. The forward mutation reporter genes previously assessed (*URA3/5, FRR1* and *UXS1*) detected only mutations that disrupt gene function, resulting in drug resistance on selective medium (31). This method was effective for detecting novel Tcn12 insertions, which, as demonstrated in this study, inserted primarily in gene-coding regions (*SI Appendix*, Fig. S7). These loss-of-function reporter systems were not, however, able to detect TE movements into non-coding regions, where we discovered movements of Cnl1 and the majority of novel T1 insertions (Fig. 4, Fig. 5, and *SI Appendix*, Table 1). In our genome-wide assessment, we found that T1 inserted disproportionately between genes. Given the small distance between most genes in Cryptococcus, insertions were often near transcription start sites (*SI Appendix*, Fig. S5). Because DNA transposons do not typically display target-specific integration (57), chromatin accessibility may influence T1 integration. For example, an assessment of integration sites by the Hermes DNA transposon introduced into *S. cerevisiae* showed that the element targeted nucleosome-free chromatin found near the 5’ ends of genes, resulting in the overrepresentation of insertions between divergent genes, with insertions between convergent genes less frequent than expected (58). We found a similar pattern of T1 integration in the passaged TA lines in XL280α, with a strong bias for insertion between divergent genes and against insertion between convergently orientated genes. Alternatively, the T1 transposase may be tethered to proteins that preferentially bind near promoter regions; this mechanism of “integrase-tethering,” however, has been described primarily for some LTR retrotransposons (57).

The insertion of Tcn12 and T1 into gene-coding regions (*SI Appendix*, Table S1) is likely to disrupt the function of the corresponding gene. However, for those T1 insertions that occurred near promoter regions, these movements could have a major impact on modulating gene expression and phenotypic outcomes of nearby genes. More global effects would be expected if the expression of a transcription factor is affected. In Cryptococcus, TE-mediated changes in gene expression could alter virulence factor production, resulting in differences in capsule or melanin production, for example. Unexplained phenotypic differences between incident and relapse *C. neoformans* isolates in patients with recurrent cryptococcal meningoencephalitis have been reported (15, 59). In these cases, there were a number of phenotypic changes observed between paired isolates from the same patient, but the genetic basis for many remains unknown. While epigenetic modifications may drive some phenotypic changes, it is also possible that TE movements undetected by traditional short-read sequencing may be responsible for others. In a 2013 study of recurrent cryptococcal infections, for example, changes in the location and copy number of the Tcn6 retrotransposon were detected by Southern analysis comparing incident and relapse cases of cryptococcal meningitis from the same patient (59). However, the contribution of Tcn6-mediated genomic changes to phenotypic variation between the isolates was not explored.

The estimated acquisition rate for genetic polymorphisms (SNPs/INDELs) in Cryptococcus during infection is very low—1 change in 58 days and 4 changes in 77 days in two studies that examined microevolution in *C. neoformans* during human infection (15, 59). In *C. deneoformans* lines passaged in vitro, we found that SNPs and INDELs accumulated at a similar rate per generation as TE mutations at 30° (Table 2). At the elevated temperature of 37°, however, the rate of TE mutations increased by five-fold, while the rate of SNPs and INDELs remained the same. Therefore, depending on the particular infecting strain of Cryptococcus and the number of mobile elements in its genome, the frequency of TE-mediated mutations in vivo may be much higher than that of minor sequence variations. Our finding that multiple TE insertions occurred in single genomes during a ten-day murine model of infection supports this hypothesis (Fig. 6). By contrast, a recent study in the plant pathogenic fungus *Zymoseptoria tritici* reported an increase in base substitutions, deletions and duplications in response to a mild temperature stress (28° compared to 18°), but did not detect an increase in the rate of insertions (60); it is unclear, however, if the mobility of TEs was specifically assessed.

Unlike T1 and Tcn12 insertions that occurred within and between genes, Cnl1 insertions into non-coding, sub-telomeric regions of chromosomes are unlikely to alter gene expression or disrupt gene function. However, as the number of genomic Cnl1 copies increases, the likelihood that the element could insert into genes or in proximity to genes also increases. This was demonstrated in a recent study investigating two clinical hypermutator strains of *C. neoformans* (33). Due to a nonsense mutation in *ZNF3* that inactivated RNAi suppression of Cnl1, the copy number in the telomeres of most chromosomes of these strains had reached ~100-150 full-length copies, with numerous additional truncated copies. In these strains harboring a heavy Cnl1 burden, the *FRR1* reporter gene was disrupted by Cnl1 insertion, resulting in rapamycin+FK506 drug resistance. It is therefore possible that accumulation of high copy numbers of Cnl1 during growth in the environment or during persistent host infections could result in Cnl1 movements that affect phenotypes or infection outcomes, particularly if Cnl1-mediated resistance to antifungal treatment is a consequence.

The mechanism(s) of heat stress-induced TE mobility in Cryptococcus remains undetermined but may be linked to global changes in transcription and translation that occur during cellular adaptation to thermal stress. A mechanism for heat stress-induced transposition was described for the *Ty1*/*copia*-like LTR retrotransposon, *ONSEN*, found in Brassicaceae plant species (24). *ONSEN* mobilizes in concert with the plant cell’s heat stress response through heat response element sequences in its LTR that are recognized by the major heat response transcription factor HsfA2. Heat shock element (HSE) sequences are conserved in eukaryotes and are recognized by global regulators of the heat shock response (61). In Cryptococcus, Hsf1 is the major transcription factor that activates genes required for survival at 37°, including *HSP90* (62). HSE sequences were identified upstream of several genes directly controlled by Hsf1 in *C. neoformans*, and most were “step-type HSEs”, consisting of three nTTCn or nGAAn pentanucleotide repeats interrupted by five bp sequences (62). In scanning the LTR of Tcn12 copies in both *C. neoformans* H99 and *C. deneoformans* XL280α, we identified a putative HSE sequence upstream of the predicted transcription start for the *gag-pol* polyprotein of Tcn12. This putative HSE (T<u>TTC</u>TTTTAGA<u>TTC</u>TACGATA<u>GTC</u>T) is imperfect by only a single base-pair in the terminal nTTCn repeat, and resembles the imperfect step-type HSEs identified in *C. neoformans* (62). Whether Hsf1 acts directly to activate Tcn12 transposition in Cryptococcus remains to be tested.

The fact that mutation accumulation lines or reporter assays are needed to detect TE movements in Cryptococcus means that these movements are infrequent. Understanding the sporadic nature of TE movements in individual cells and identifying those stimuli that trigger movements are important and active areas of investigation. In the human host, where cryptococci can persist for weeks or months to even years and decades, there is a strong selective pressure for cells that evolve adaptive traits. Mammalian core body temperatures currently pose a significant stress for the majority of environmental fungi, limiting the number of species capable of causing human infection and disease (10). In light of rising global temperatures, however, environmental fungi (both pathogenic and benign) are likely to evolve increased thermotolerance. In addition, greater fungal spore dispersal due to climate change and habitat disruption is predicted to bring more fungal diseases to the human population (63, 64). Heat stress-induced genetic changes, including increased TE mutations, suggest that pathogenic traits in environmental fungi may evolve more rapidly than anticipated, increasing the urgency to develop antifungal vaccines and to augment our limited arsenal of antifungal drugs.

## Materials and Methods

### Strains and growth media

The *C. deneoformans* XL280α wildtype strain (65, 66) and the *RDP1* mutant of this strain (*rdp1Δ∷NAT*) (31) served as the parental lineages for all mutation accumulation lines generated. Strains were grown in non-selective YPD medium (1% yeast extract, 2% Bacto Peptone, 2% dextrose, and 2% agar for plates) at 30° for all experiments except for TA lines maintained at 37° during passage.

### Mutation accumulation lines

TA lines were generated by single colony, bottleneck passage 40 times onto YPD agar. Single colonies from each TA line were selected randomly for passage and re-streaked onto fresh medium after two days incubation at 30° or three days at 37°, when colony sizes were similar. To confirm that a similar number of generations occurred at each passage, the number of generations of growth was approximated by determining the cell count of three independent colonies at the time of passage for each incubation period at 30° or 37°. Glycerol stocks were prepared of each TA line every ten passages for most lines or at least twice during passage.

### Southern analysis

Genomic DNA was prepared using a CTAB-DNA extraction method (67), digested with restriction enzymes (New England Biolabs) and transferred to a nylon membrane (Roche). Target probes were prepared using chemiluminescent DIG-labeling (Roche); see *SI Appendix*, Table S4 for primers used. Hybridization and detection (Amersham Hyperfilm ECL) were performed according to recommended manufacturer protocols.

### Quantitative RT-PCR

RNA was extracted from mid-log phase cultures using the TRIzol reagent and protocol (Ambion). RNA was treated with DNase using the DNA-free kit (Invitrogen) and cDNA prepared using the High Capacity RNA-to-cDNA Kit (Applied Biosystems). Quantitative PCR was performed using QuantStudio Pro real-time PCR system with Power Track SYBR Green Master Mix reagents (Applied Biosystems).

### Whole-genome sequencing, TE mapping, variant calling, 5mC methylation detection

Provided in the Supplementary Information text.

## Data Availability

Paired-end, Illumina sequence reads, as well as Nanopore sequence reads, from TA lines are hosted on NCBI’s SRA under BioProject PRJNA794155, with accession numbers SRR17430287 – SRR17430309 and SRR17722984 – SRR17722990, respectively. Python scripts used in the analysis of TE movement and accumulation (and figure generation) in TA lines are hosted at: https://github.com/magwenelab/Transposon-mobility

## Acknowledgments

We would like to thank Márcia David-Palma for her training assistance with Nanopore sequencing. This research was supported by National Institutes of Health grants R35GM118077 and R21AI133644 to S.J.R, the NIH Tri-Institutional Molecular Mycology and Pathogenesis Training Program postdoctoral fellowship (5T32AI052080 to J.D.W and 2T32AI052080 to A.G.), and the NIH/NIAID K99 grant 1K99AI166094-01 to A.G. Research was also supported in part by NIH/NIAID R01 grants AI039115-24 and AI050113-17 to J.H., and AI133654-05 to J.H. and P.M.

