## Supplementary Information for "Genome-wide analysis of heat stress-stimulated transposon mobility in the human fungal pathogen *Cryptococcus deneoformans*"

*Sue Jinks-Robertson

**This PDF file includes:**

Supplementary text
Figures S1 to S11
Tables S1 to S5
SI References

### Supplementary Information Text

### Materials and Methods

#### Illumina sequencing and TE mapping

Pair-end reads from whole-genome sequencing were analyzed to identify discordant pairs mapping between a TE and the XL280α reference genome (similar to analysis presented in (1)). This was done by first identifying reads that align to the sequence of a TE (either the T1 or Tcn12 element) with BLAT – the BLAST like alignment tool (2) – using default settings. The mated pairs of these TE-associated reads that failed to align to the TE were then mapped (again using BLAT) to the XL280α genome. Reads mapping to the TE were labeled as supporting reads while their mates that mapped to the reference genome were denoted as anchor reads. Anchor reads were only considered if they mapped uniquely to the XL280α genome. Per chromosome, windows of 10 kb were used to quantify and identify the number and position of anchor reads throughout the genome. This analysis was conducted on both TA lines and the non-passaged XL280α strain.

Two methods of standardization were used to identify TE insertion and copy loss in TA lines, using the corresponding signal of anchor reads in non-passaged strains. For TE insertions in TA lines, the number of anchor reads (per 10 kb) was divided by the corresponding number of TE anchor reads (plus one) within the non-passaged strain. For TE copy loss, the anchor read signal of each non-passaged strain was divided by the negative of the anchor read count (plus one) of the given TA line. When plotted, this generates a positive or negative, genome-wide signal of anchor reads relative to the non-passaged strain for TE insertions or loss (respectively) in each TA line.

Loci that demonstrated a potential insertion or loss of a TE were identified per TA line by retaining 10 kb windows with greater than 10x (+ or -, respectively) anchor read coverage, removing regions that were identified in all TA lines, and regions with a position in a native TE site of the XL280α reference genome. The locations of predicted TE insertions and TE copy losses in passaged TA lines were confirmed by PCR and Sanger sequencing (see *SI Appendix,* Table S4 for primers used).

**Variant Calling**

Pair-end Illumina-sequenced reads for each sample were aligned to the nanopore XL280α reference genome using bwa-mem (3) and its default settings. Duplicate reads were identified using samblaster (4). Duplicate marked alignments were filtered via the samtools view command (5) with the additional filtering options of -F 3852 and mapping quality threshold of ten (with -q 10). Read groups were added to each binary alignment map file using bamaddrg (6).

All filtered alignments were fed into samtools mpileup and bcftools to identify genetic variants segregating across the passaged isolates. The default parameters were used in the call to mpileup with the addition of the -d 100 flag to cap the maximum number of aligned reads considered in detecting variants (6). The default consensus calling strategy in bcftools, with a ploidy setting of one and the addition of the options "-vc" were used to detect variants.

Variant calls were filtered using bcftools and custom scripts in python. Specifically, using the bcftools view command, only variants with a quality score greater than 900 were considered. Invariant sites – loci with fixed genetic variants across all samples – were removed from analysis. Per isolate, variant loci were considered only if they had greater than 6x coverage with 80% of reads suggesting a variant call. Per isolate, only variants detected and consistent across each technical replicate were retained for further consideration.

Using the Integrative Genomics Viewer (IGV), variant positions were visually inspected to remove those called within telomeric regions that displayed as artifacts, i.e. appearing with identical alignments across multiple, unrelated isolates in the passaging experiments (7). In total, 836 raw genetic variants were called via bcftools and after scripted filtering, 198 variants were examined in IGV, resulting in the identification and analysis of 142 unique variant sites. IGV was also used to examine inheritance patterns of genetic variants and estimate potential changes in amino acid sequences across isolates.

**Nanopore sequencing and TE mapping**

High molecular-weight DNA was isolated by a CTAB-DNA extraction method (8) and tested for its quality using NanoDrop. Samples were sequenced on the MinION system using the R9.4.1 Flow-Cell and SQK-LSK109 library preparation kit. For multiplexing of multiple samples in a single flow-cell, Native Barcoding Expansion 1-12 kit (EXP-NBD104) was used during the library preparation as instructed by the manufacturer's protocol. Nanopore sequencing was performed at the default voltage for 48 hours as per the MinION sequencing protocol provided by the manufacturer. MinION sequencing protocol and setup was controlled using the MinKNOW software. Base-calling was performed outside the MinKNOW software on the lab server using the standalone Guppy basecaller from Nanopore and the sequence reads thus obtained were used for genome assembly. In case of multiplexed runs, the reads were de-multiplexed using qcat or Guppy_barcoder before the genome assembly. Canu (9) was used to assemble the genome of most strains using reads that are longer than 2 kb (-minReadLength=2000) in length. The genome assembly was checked for integrity by mapping the reads back to the genome assembly using minimap2 (10), and duplicated small contigs were discarded. The contigs that were broken at centromeres were assembled manually after a synteny comparison and analysis using Geneious Prime 2020.1.1. All the chromosome-level contigs thus generated were then reoriented to have the same configuration as the wildtype XL280α genome. Transposon locations were determined using BLASTn analysis and annotated using Geneious Prime 2020.1.1.

**Detection of 5mC methylation**

5mC DNA methylation was calculated using the scripts available with nanopolish, as previously described (11). Briefly, the XL280α nanopore reads were mapped to the genome assembly, and methylation status was identified using "call-methylation" followed by calculating methylation frequency. The frequency was then converted to bedGraph for visualization purposes in IGV and the final figures were generated in Adobe Illustrator.

**Gene liftover and orientation analysis**

Individual gene sequences were taken from the JEC21 reference genome (version 48, FungiDB) and mapped via BLAT to the assembled XL280α nanopore reference genome.
After this mapping, those gene sequences with greater than 90% sequence identity were retained for analysis. Per chromosome, the distance between and orientation of pairs of genes were determined via python.

### SI Appendix Figures

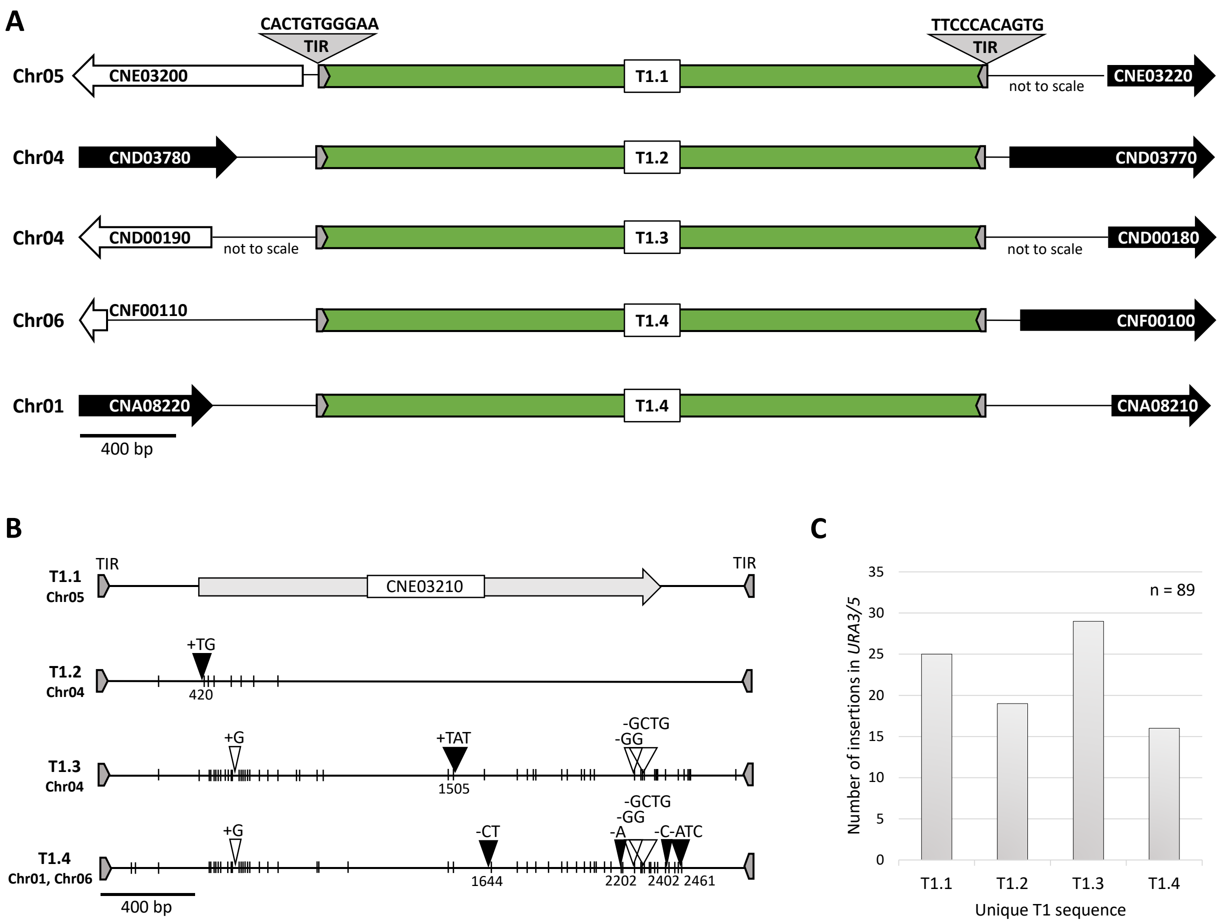


**Figure S1.** Locations of the four unique 2.7 kb T1 mobile elements in the XL280α genome. *(A)* Gene-level view of T1 copies and proximity to nearby genes. The sequence of the 11-bp terminal inverted repeats (TIRs) is indicated. *(B)* Sequence differences that distinguish each T1 element from T1.1. Single lines indicate SNPs, triangles indicate INDELs; black triangles indicate INDELs diagnostic for each element. *(C)* Number of *de novo* insertions of each T1 element in the *URA3/5* genes of 5FOA-resistant mutants (12).

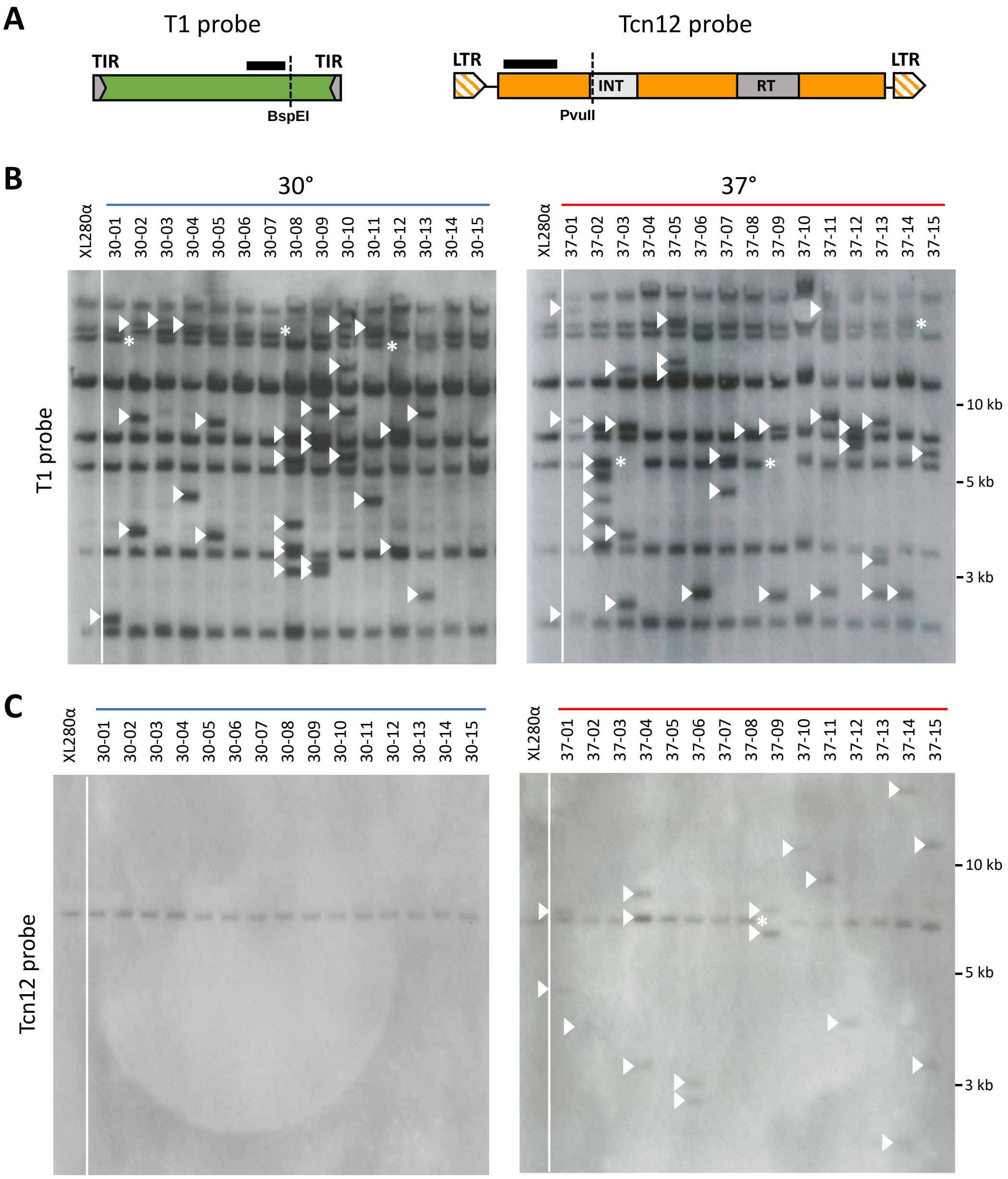


**Figure S2.** Southern analysis of TE movement in fifteen independent XL280α TA lines passaged ~800 generations on YPD at 30° or 37° as indicated. *(A)* Location of the restriction sites (dotted lines) and probes (black bars) used in Southern analysis. *(B)* Genomic DNA of non-passaged XL280α (left lane) and passaged isolates digested with BspEI, probed for T1. *(C)* DNA digested with PvuII, probed for Tcn12. Arrowheads indicate *de novo* TE insertions and asterisks indicate the loss of a native TE copy.

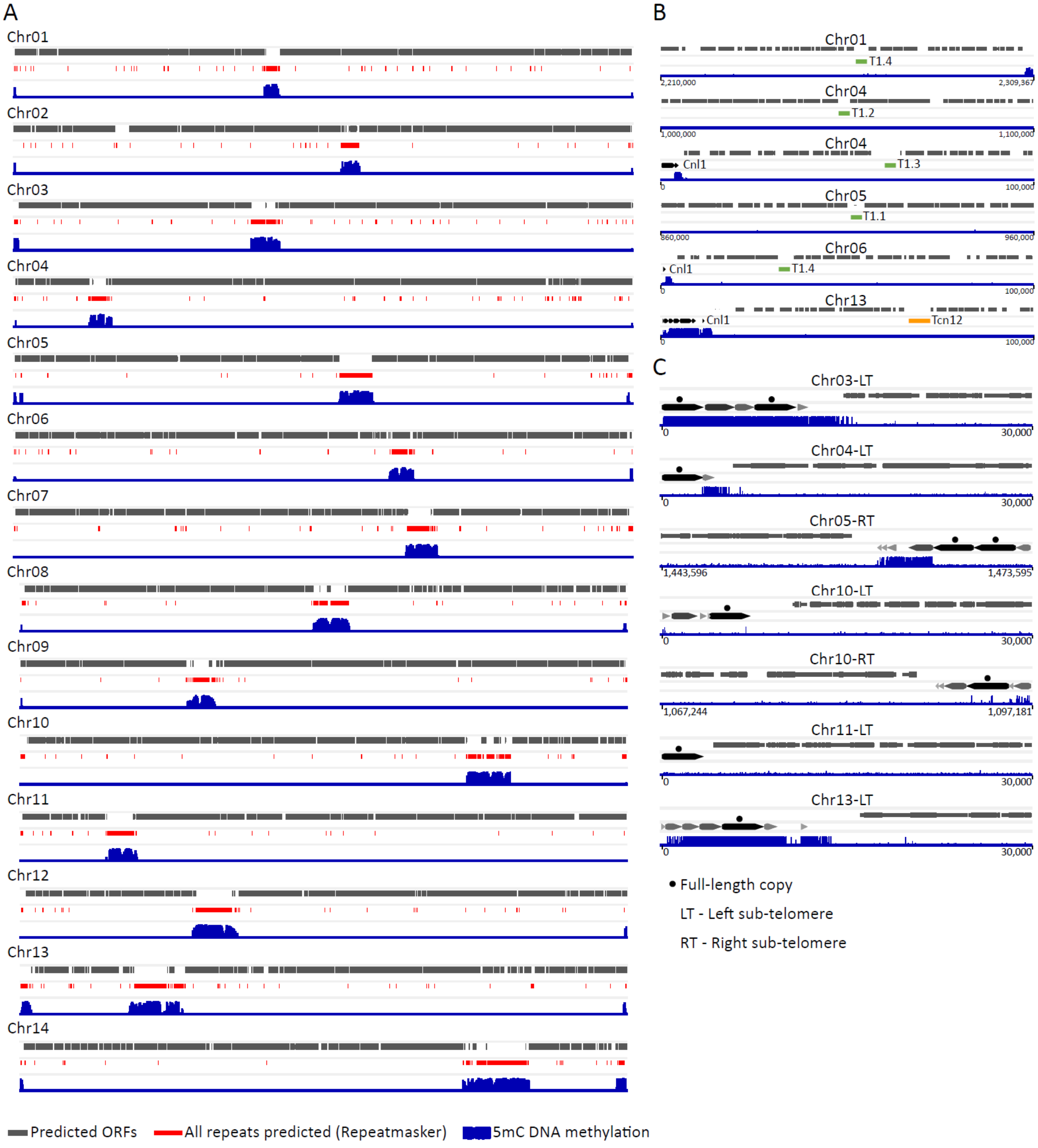


**Figure S3.** 5-methylcytosine (5mC) DNA methylation in XL280α as detected by Nanopore sequencing. *(A)* Genome-wide 5mC methylation pattern with predicted open reading frames (ORFs) and repetitive element sequences. 5mC methylation at regions of chromosomes encoding mobile T1 and Tcn12 sequences *(B)* and Cnl1 sequences, zoomed in to show full-length copies (black arrowheads) and truncated copies (gray arrowheads) in the left and right sub-telomeres *(C).*

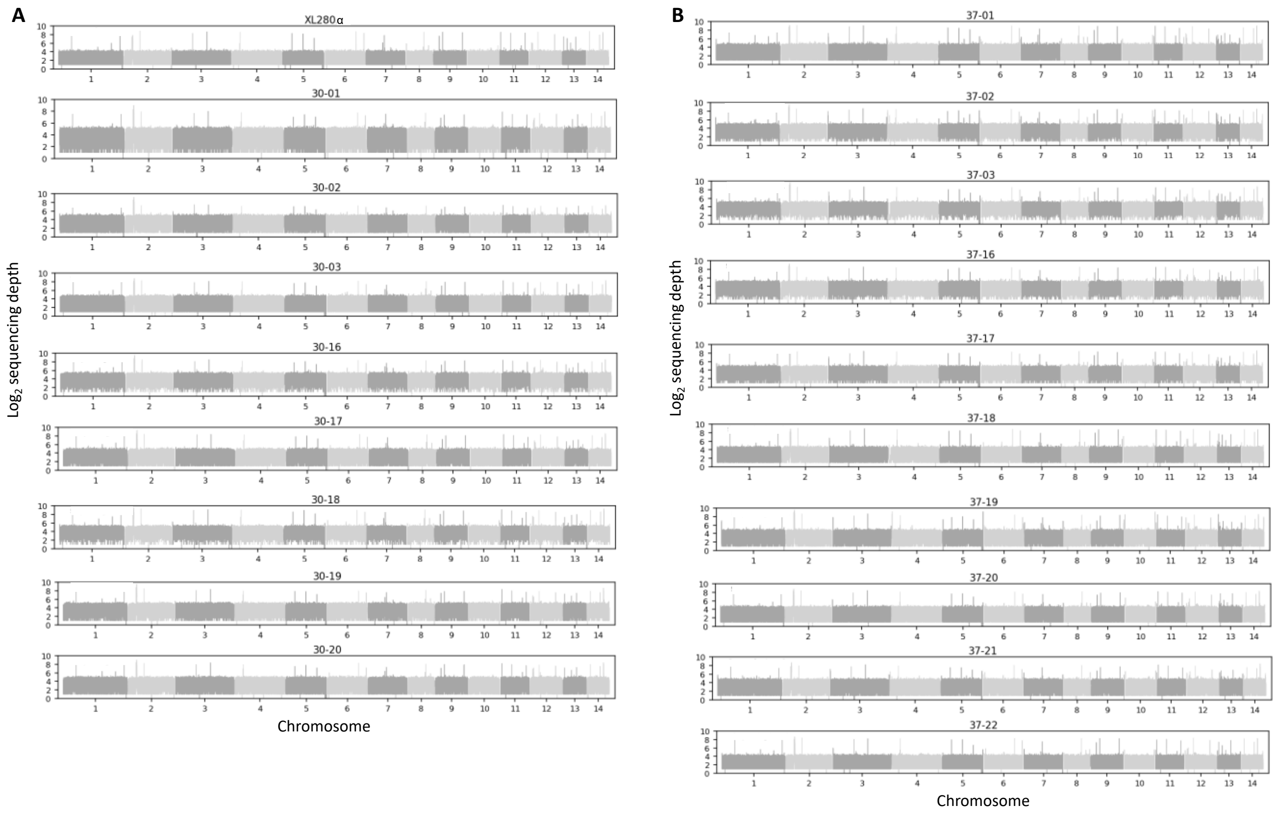


**Figure S4.** Sequencing depth of Illumina reads for the genomes of *(A)* XL280α and TA lines passaged at 30° and *(B)* TA lines passaged at 37°. Normalized read counts were transformed to the $log_2$ scale for sequencing depth.

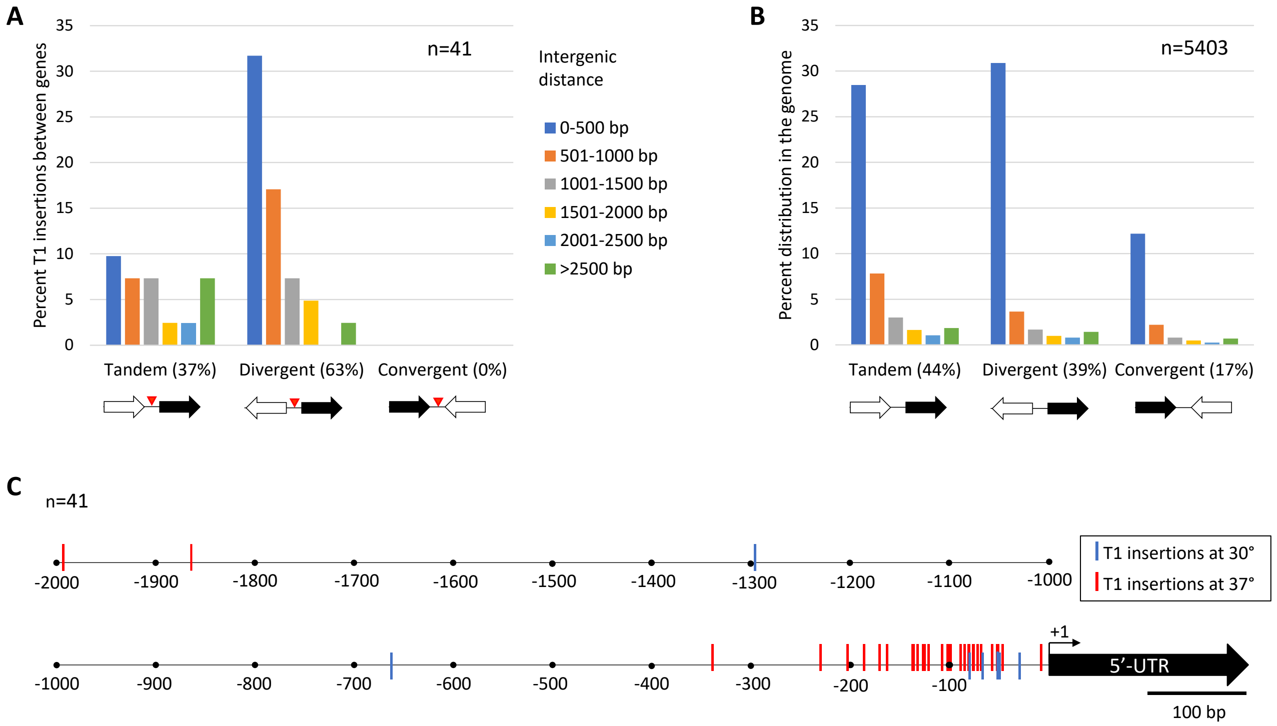


**Figure S5.** Analysis of *de novo* T1 insertions between genes in XL280α TA lines. *(A)* Percentage of mapped T1 insertions (red arrowheads) between genes in tandem, divergent or convergent orientations. *(B)* Distribution of tandem, divergent and convergent gene orientations in the XL280α genome. Overlapping genes in all orientations were excluded. *(C)* Distance of mapped T1 insertions upstream of the nearest +1 start of transcription (as annotated for the *C. deneoformans* JEC21 reference genome, EnsemblFungi).

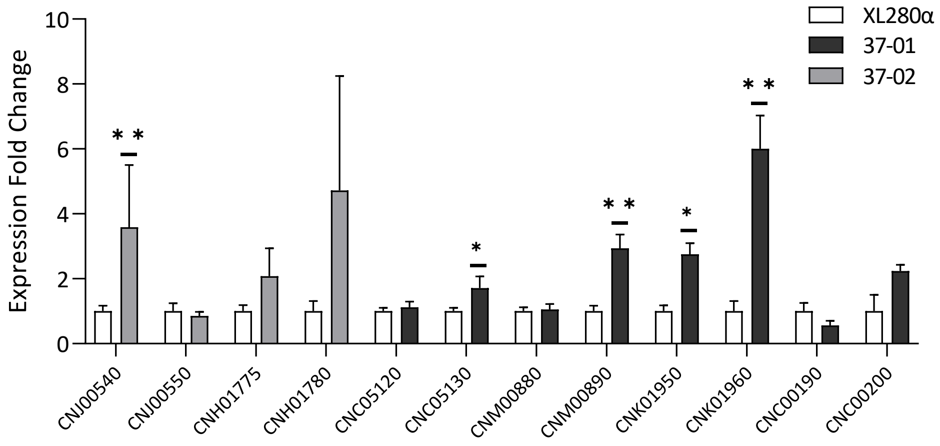


**Figure S6.** Fold change in expression of control genes in 37°-passaged lines 37-01 and 37-02 relative to the parental XL280α strain. These control genes are not proximal to the *de novo* T1 insertions in 37-01 and 37-02 shown in Fig 4. Gene expression levels were normalized to the endogenous reference gene *GPD1* using the comparative $\Delta\Delta C_T$ method. For each target gene and each sample, technical triplicate and biological triplicate qRT-PCR reactions were performed. Error bars represent standard error of the means (SEM) for three biological replicates. Statistical analyses were performed using the GraphPad Prism 9 software. A Welch's unpaired t test (two-tailed) was performed for each pairwise comparison using the mean $\Delta C_T$ values for three biological replicates (ns, not significant; * indicates $0.01<p\leq0.05$; and ** indicates $0.001<p\leq0.01$).

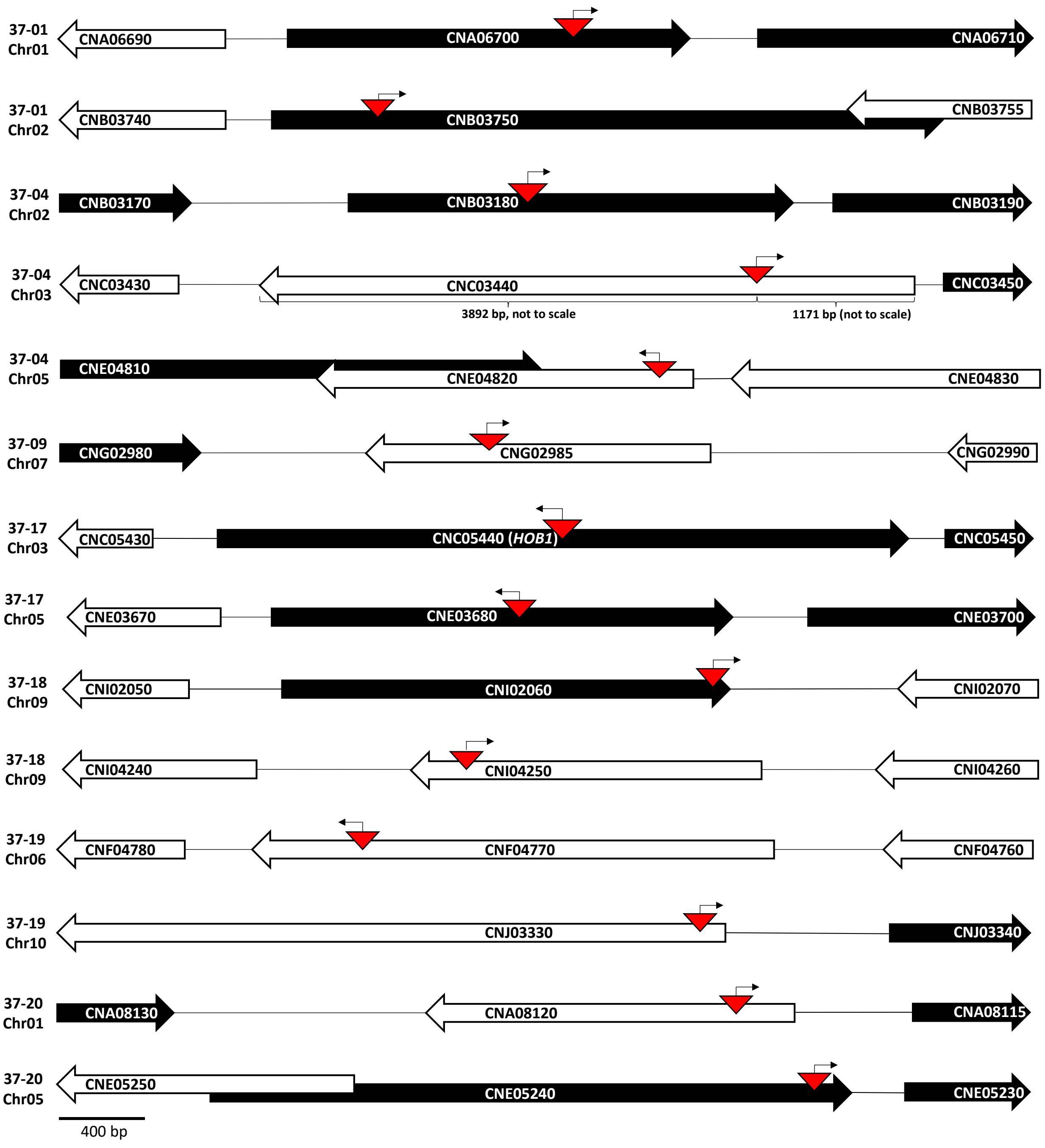


**Figure S7.** Analysis of *de novo* Tcn12 integration sites in TA lines passaged at 37°. Gene-level view of the 14 novel Tcn12 insertions (triangles) within gene-coding regions. Arrows indicate the forward or reverse orientation of Tcn12 with respect to the *gag-pol* polyprotein encoded within the element.

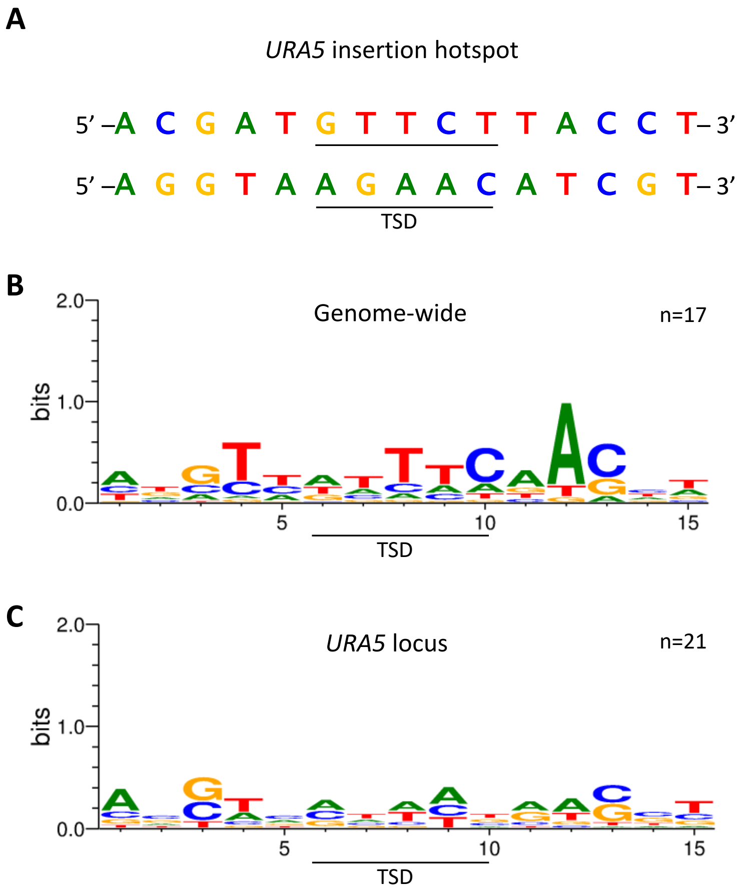


**Figure S8.** Alignment of *de novo* Tcn12 integration sites with respect to the forward orientation of the element. *(A) URA5* insertion hotspot for Tcn12, centered at the TSD [6]. The 15-bp 5’ to 3’ *URA5* sequence is shown for both DNA strands. *(B)* Sequence logo (WebLogo 3.7.4) generated by aligning 17 *de novo* Tcn12 integration sites (Table S3) in 37° TA lines. *(C)* Sequence logo generated by aligning unique Tcn12 integration sites in the *URA5* locus of 5FOA-resistant XL280α mutants recovered from mouse organs [6].

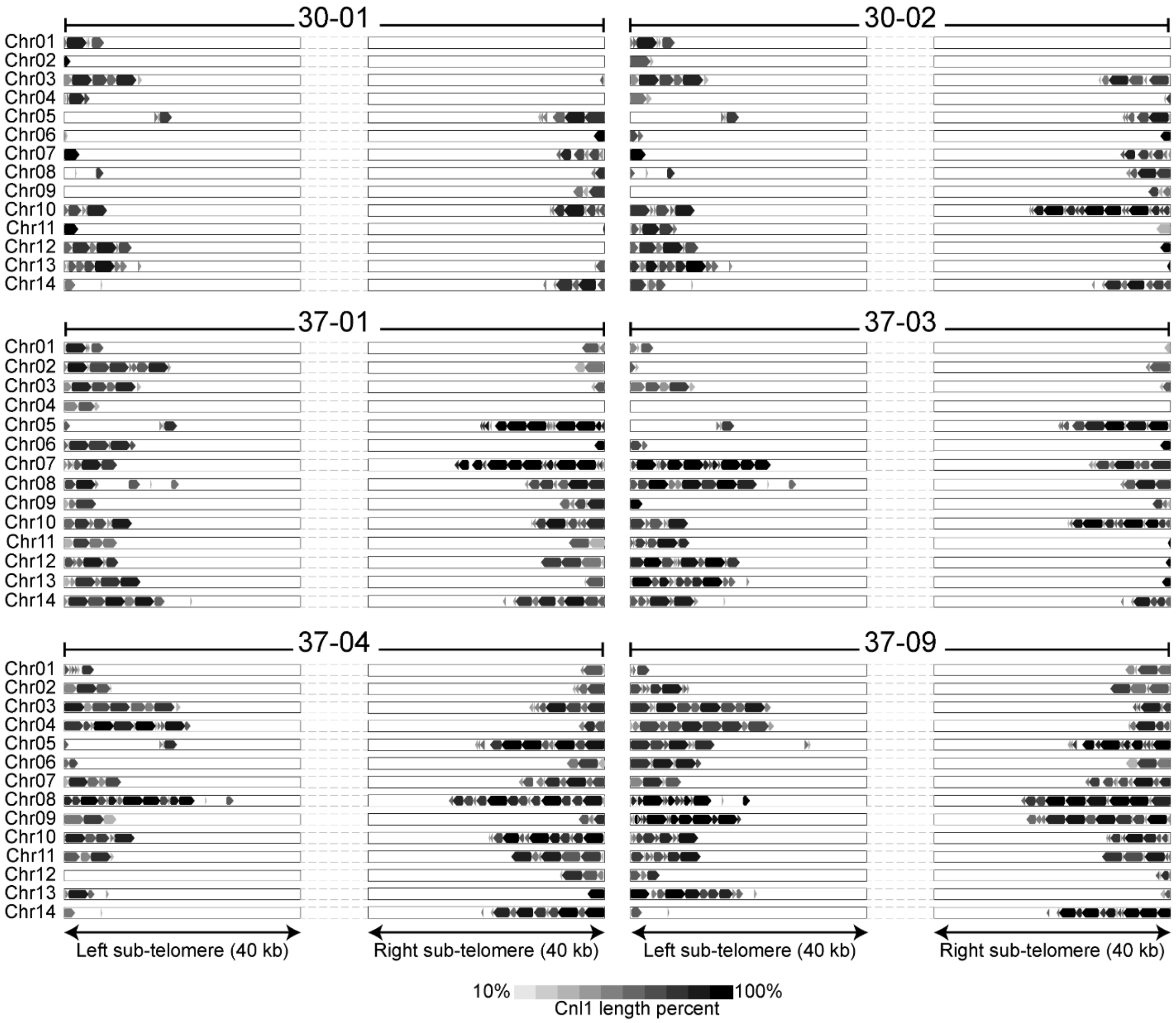


**Figure S9.** Full-length and truncated copies of the non-LTR Cnl1 retrotransposon at the sub-telomeres of chromosomes of TA lines passaged at 30° (30-01, 30-02) and 37° (remaining isolates), sequenced by Nanopore. Left and right panels for each isolate indicate the left or right sub-telomeric region as indicated. The size of each block arrows approximates the percent length of the 3.4 kb Cnl1 element with the darkest color (black) indicating full-length copies and lighter shades of gray indicating truncated copies of the element. The distribution of Cnl1 on chromosomes for the non-passaged XL280α parent is shown in Fig. 5A.

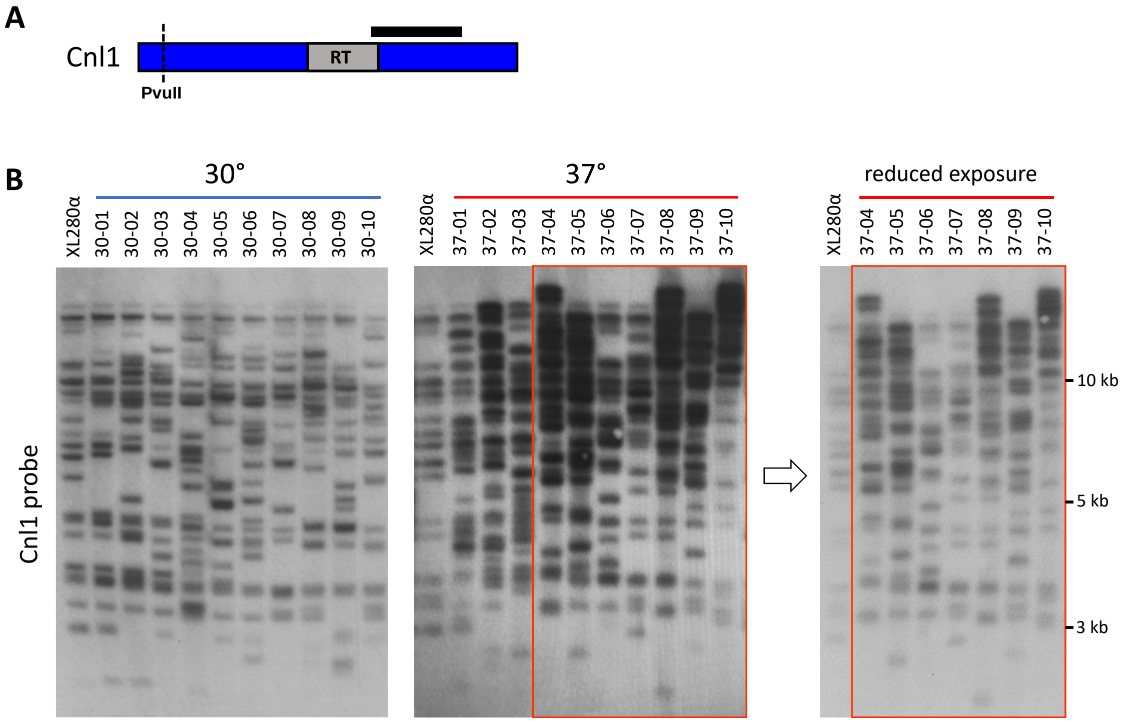


**Figure S10.** Southern analysis probing for the Cnl1 retrotransposon in the genomes of XL280α TA lines passaged at 30° and 37°. *(A)* Location of the PvuII restriction site (dotted line) and probe (black bar) used to identify Cnl1 copies. *(B)* Southern analysis of Cnl1 copies in XL280α and TA lines passaged at 30° (left) or 37° (right). The far-right panel shows a reduced exposure of the 37° passaged lines indicated in the red box for better resolution of larger bands.

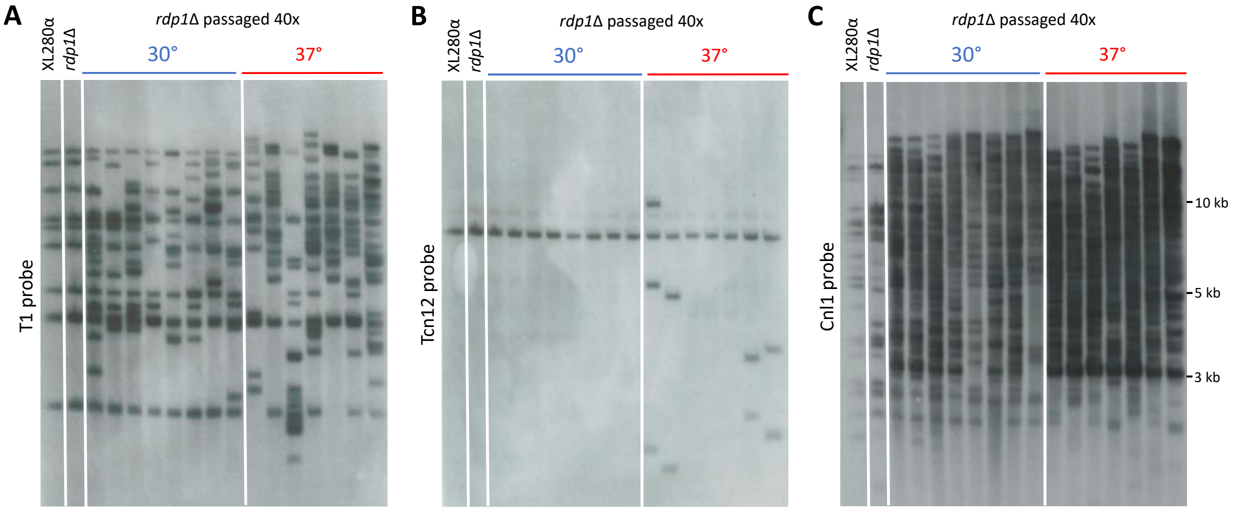


**Figure S11.** Southern analysis of TE movements in *Cryptococcus* XL280α *rdp1*Δ mutants passaged 40 times (~800 generations) at 30° and 37°. Genomic DNA from the wildtype XL280α strain, *rdp1*Δ mutant strain and passaged *rdp1*Δ TA lines was digested with PvuII and probed for *(A)* T1, *(B)* Tcn12, and *(C)* Cnl1.

**Table S1.** Novel T1 and Tcn12 insertions mapped in lines passaged 40 times at 30° or 37°. Genomic locations of novel TEs are defined as upstream (US) or downstream (DS) of predicted transcription start sites for proximal genes, as annotated for the *C. deneoformans* JEC21 reference genome (EnsemblFungi and FungiDB).

| Isolate | TE | Chr | Category | Location |
|---|---|---|---|---|
| 30-01 | T1 | 8 | intergenic - divergent | 51 bp US CNH02260, 109 bp US CNH02250 |
| 30-02 | T1 | 2 | intergenic - divergent | 50 bp US CNB02630, 959 bp US CNB02620 |
| 30-02 | T1 | 3 | intragenic | 4484 bp into CNC05220 |
| 30-02 | T1 | 11 | intergenic - divergent | 50 bp US CNK03170, 1014 bp US CNK03180 |
| 30-02 | T1 | 13 | intergenic - divergent | 665 bp US CNM02240, 1137 bp US CNM02230 |
| 30-03 | T1 | 7 | intergenic - divergent | 66 bp US CNG03170, 115 bp US CNG03180 |
| 30-16 | T1 | 3 | intergenic - tandem | 1294 bp US CNC01660, 640 bp DS CNC01670 |
| 30-16 | T1 | 6 | intragenic | 3115 bp into CNF02870 |
| 30-17 | T1 | 6 | intergenic - divergent | 80 bp US CNF00080, 93 bp US CNF00090 |
| 30-20 | T1 | 10 | intergenic - divergent | 28 bp US CNJ02860, 501 bp US CNJ02870 |
| 37-01 | T1 | 7 | intergenic - divergent | 340 bp US CNG02000, 972 bp US CNG01990 |
| 37-01 | T1 | 8 | intragenic | 40 bp into CNH01775 (5'-UTR), 99 bp US CNH01780 |
| 37-01 | T1 | 10 | intergenic - divergent | 52 bp US CNJ00550, 146 bp US CNJ00540 (*LYS2*) |
| 37-01 | T1 | 10 | intragenic | 195 bp into CNJ02670 (5'-UTR), 1526 bp DS CNJ02660 |
| 37-01 | Tcn12 | 1 | intragenic | 1300 bp into CNA06700 |
| 37-01 | Tcn12 | 2 | intragenic | 480 bp into CNB03750 |
| 37-01 | Tcn12 | 8 | intragenic | within Cnl1 left arm; subtelomere |
| 37-02 | T1 | 1 | intergenic - divergent | 99 bp US CNA06060, 491 bp US CNA06050 |
| 37-02 | T1 | 1 | intergenic - tandem | 1866 bp US CNA07370, 274 bp DS CNA07360 |
| 37-02 | T1 | 2 | intergenic - tandem | 170 bp US CNB02680, 3813 bp DS CNB02690 |
| 37-02 | T1 | 3 | intergenic - divergent | 102 bp US CNC00190, 144 bp US CNC00200 |
| 37-02 | T1 | 3 | intergenic - divergent | 89 bp US CNC05130, 90 bp US CNC05120 |
| 37-02 | T1 | 4 | intergenic - tandem | 232 bp US CND03560, 790 bp DS CND03550 |
| 37-02 | T1 | 4 | intergenic - tandem | within a tRNA (CND05420); 203 bp US CND05430, 118 bp DS CND05410 |
| 37-02 | T1 | 11 | intergenic - divergent | 76 bp US CNK01950, 105 bp US CNK01960 |
| 37-02 | T1 | 13 | intergenic - divergent | 50 bp US CNM00890, 132 bp US CNM00880 |
| 37-02 | Tcn12 | 2 | other | centromere, near CNB02910 |
| 37-03 | T1 | 2 | intragenic | 1212 bp into CNB01810, 106 bp US CNB01800 (*FKBP12*) |
| 37-03 | T1 | 2 | intergenic - divergent | 135 bp US CNB05060, 682 bp US CNB05080 |
| 37-03 | T1 | 3 | intergenic - tandem | 71 bp US CNC00160, 584 bp DS CNC00170 |
| 37-03 | T1 | 4 | intergenic - tandem | 132 bp US CND05850, 272 bp DS CND05860 |
| 37-03 | T1 | 6 | intergenic - divergent | directly US of native T1.4 copy; 866 bp DS of CNF00110 |
| 37-03 | T1 | 11 | intergenic - divergent | 106 bp US CNK01520, 110 bp US CNK01510 |
| 37-04 | Tcn12 | 2 | intragenic | 830 bp into CNB03180 |
| 37-04 | Tcn12 | 3 | intragenic | 3892 bp into CNC03440 |
| 37-04 | Tcn12 | 5 | intragenic | 155 bp into CNE04820 |
| 37-04 | Tcn12 | 12 | other | telomere region |
| 37-09 | T1 | 6 | intergenic - divergent | 66 bp US of CNF00100, 60 bp DS of native T1.4 copy |
| 37-09 | T1 | 10 | intergenic - divergent | 100 bp US CNJ00250, 552 bp US CNJ00260 |
| 37-09 | Tcn12 | 7 | intragenic | 1032 bp into CNG02985 |
| 37-09 | Tcn12 | 13 | other | non-coding region near CNM00220 |
| 37-16 | T1 | 7 | intergenic - divergent | 84 bp US CNG00580, 1504 bp US CNG00590 |
| 37-17 | Tcn12 | 3 | intragenic | 1569 bp into CNC05440 |
| 37-17 | Tcn12 | 5 | intragenic | 1170 bp into CNE03680 |
| 37-18 | T1 | 2 | intergenic - tandem | 1995 bp US of CNB02910, 647 bp DS CNB02900 |
| 37-18 | T1 | 14 | intergenic - tandem | 188 bp US CNN00920, 68 bp DS CNN00910 |
| 37-18 | Tcn12 | 9 | intragenic | 1374 bp into CNI04250 |
| 37-18 | Tcn12 | 9 | intragenic | 1982 bp into CNI02060 |
| 37-19 | T1 | 1 | intergenic - tandem | 162 bp US CNA01370, 709 bp DS CNA01360 |
| 37-19 | T1 | 1 | intergenic - tandem | 52 bp US CNA08100, 384 bp DS CNA08090 |
| 37-19 | T1 | 6 | intergenic - tandem | 46 bp US CNF01870, 476 bp DS CNF01880 |
| 37-19 | T1 | 13 | intergenic - divergent | 47 bp US CNM02270, 448 bp US CNM02280 |
| 37-19 | Tcn12 | 6 | intragenic | 1917 bp into CNF04770 |
| 37-19 | Tcn12 | 10 | intragenic | 191 bp into CNJ03330 |
| 37-20 | T1 | 1 | intergenic - tandem | 122 bp US CNA03220, 3027 bp DS CNA03240 |
| 37-20 | Tcn12 | 1 | intragenic | 267 bp into CNA08120 |
| 37-20 | Tcn12 | 5 | intragenic | 2791 bp into CNE05240 |
| 37-21 | T1 | 4 | intergenic - divergent | 57 bp US CND00300, 3545 bp US CND00310 |
| 37-21 | T1 | 4 | intergenic - divergent | 80 bp US CND03930, 396 bp US CND03920 |
| 37-21 | T1 | 9 | intergenic - divergent | 135 bp US CNI01880, 144 bp US CNI01890 |
| 37-21 | Tcn12 | 4 | intergenic | 410 bp US CND04840 |
| 37-22 | T1 | 1 | intergenic - tandem | 6 bp US CNA00670, 1407 bp DS CNA00660 |
| 37-22 | T1 | 2 | intergenic - tandem | 76 bp US CNB02550, 1081 bp DS CNB02545 |
| 37-22 | T1 | 3 | intergenic - divergent | 68 bp US CNC04820, 187 bp US CNC04810 |
| 37-22 | T1 | 5 | intergenic - divergent | 82 bp US CNE03320, 532 bp US CNE03310 |

**Table S2.** Distribution of genes in tandem, divergent and convergent orientation in the XL280α genome compared to the distribution of novel T1 insertions between genes in each orientation. Overlapping genes in all orientations were excluded, and predicted genes based on those annotated in the JEC21 reference genome (EnsemblFungi).

| Gene orientation | XL280α genome | | Intergenic T1 insertions | | Difference |
|---|---|---|---|---|---|
| | Number | Percent | Number | Percent | p-value |
| Tandem | 2371 | 44% | 15 | 37% | p=0.43 |
| Divergent | 2133 | 39% | 26 | 63% | p=0.0030* |
| Convergent | 899 | 17% | 0 | 0% | p=0.0081* |
| TOTAL | 5403 | 100% | 41 | 100% | |

*statistically significant (p<0.017) by Chi-square test of association applying the Bonferroni correction. Chi-square goodness of fit global test (p=0.001).

**Table S3.** Location and alignment of novel Tcn12 insertion sites centered around the target site duplication (TSD) sequence (red letters).

| DNA sequences at novel Tcn12 insertion sites in 40x passaged isolates (37°) | | | |
|---|---|---|---|
| **Passaged Isolate** | **Chr** | **Gene** | **5'-> 3' sequence with respect to forward orientation of Tcn12** |
| 37-01 | 1 | CNA06700 | GACGGTAACATCAAATTCG |
| 37-01 | 2 | CNB03750 | GACTGTTACTTTTACTACC |
| 37-01 | 8 | Cnl1 | TCTCGCATTTCCAACCCGA |
| 37-02 | 2 | CNB02910 | CCAAGATGTTTCAACAAAG |
| 37-04 | 2 | CNB03180 | TTAACTCATTACAAGTCAC |
| 37-04 | 3 | CNC03440 | CAACCCAAAATCAAACTAT |
| 37-04 | 5 | CNE04820 | GTATGTCTTCTAAACAACC |
| 37-09 | 7 | CNG02985 | AGATCTTGTTCCCAGGTAC |
| 37-09 | 13 | non-coding | GATGGTTATTTCGGGGGCA |
| 37-17 | 3 | CNC05440 | ACTATCTAACACTACCTAG |
| 37-17 | 5 | CNE03680 | TCTTGTGACTTTTTCCATA |
| 37-18 | 9 | CNI04250 | TTACCTCTTTCCATCTTCT |
| 37-18 | 9 | CNI02060 | CGAGAGTGGTAAGAGATTG |
| 37-19 | 6 | CNF04770 | GACGGTCCAGAAGACTACC |
| 37-19 | 10 | CNJ03330 | TTGGGTCTTCGCAAGCGGT |
| 37-20 | 1 | CNA08120 | ATCTACATATTCAACAGTA |
| 37-20 | 5 | CNE05240 | ACATATTGGTTGCACGTCT |

**Table S4.** Primers used in this study.

| ID | 5'-> 3' DNA Sequence | Target region | Chr | TA line | TE |
|---|---|---|---|---|---|
| ES90 | AAGTAGCTTTCTCCTCTATCGTCCCTCCC | | | | |
| ES91 | CCTCTGTTCTTTGTGTTCGGCAATTCTCG | CNE03200/CNE03210 | 5 | XL280α | T1.1 (native) |
| ES92 | CAGGTGGACATCATCATCCAGAGGTATGC | | | | |
| ES93 | TCTTTAAAGGCATCTTGTTCCCGCTTTGC | CND03770/CND03780 | 4 | XL280α | T1.2 (native) |
| ES94 | GAGATGAACGTAGCGCGCTTAGTAGTAGG | | | | |
| ES95 | GGCTGTAAAGTAGCGAAGAAACAAAGCCC | CND00180/CND00190 | 4 | XL280α | T1.3 (native) |
| ES96 | AGTGTCGTGAGAGTAGTGACAATCAAGCG | | | | |
| ES97 | TGGGATTGAATGTGGCAGAATAGGGATCG | CNF00100/CNF00110 | 6 | XL280α | T1.4 (native) |
| ES102 | TGCCATTTTGAAGAAGAGCTGAATGGTGC | | | | |
| ES103 | AGCTCTAAGCACAGATATGTCCGACATGC | CNA08210/CNA08220 | 1 | XL280α | T1.4 (native) |
| ES62 | GTGGACGATGGTAAAACGCATTTGGTAGC | | | | |
| ES63 | CTCCGTATCCCATAGACTGGTTTTTGGGG | CNH02250/CNH02260 | 8 | 30-01 | T1 |
| ES203 | ATGAGAAAGCGTCCCGGGGACAAG | | | | |
| ES204 | TGGTAAATAAATGGAGCGAAGAGAGC | CNB02620/CNB02630 | 2 | 30-02 | T1 |
| ES134 | AAGACTGCGATGACGATGAACGCG | | | | |
| ES135 | TGGTCGACGTTACCCCATCCCATC | CNC05220 | 3 | 30-02 | T1 |
| ES136 | TCGATGATCAGAAGGGGCTCTGGC | | | | |
| ES137 | ATGGTGGTGGTCAAAGCTGTGCAC | CNK03170/CNK03180 | 11 | 30-02 | T1 |
| ES138 | CAATTTGATCCCGTGGTGGAGGGC | | | | |
| ES139 | TCCAAACCCAGCACCCAGAGACTC | CNM02230/CNM02240 | 13 | 30-02 | T1 |
| ES146 | CGCTACAGCCTTGCAGCATCTCTG | | | | |
| ES147 | CAACTGGCTGGACACAGAACTGCC | CNG03170/CNG03180 | 7 | 30-03 | T1 |
| ES140 | TCCATCCTTCCGGGCCAGTACAAG | | | | |
| ES141 | CCCGCCCAATCACAGCAATAGACG | CNC01660/CNC01670 | 3 | 30-16 | T1 |
| ES142 | GAGTCGGTGTTGACAGCAAGCTGG | | | | |
| ES143 | TTGGAGAGGGGGTCATCTTGCTCG | CNF02870/CNF02875 | 6 | 30-16 | T1 |
| ES144 | TATTCGGAGAGCTGGGTGCGGTAC | | | | |
| ES145 | AGGGTCTGATCGGGCTGGATCAAC | CNF00090/CNF00080 | 6 | 30-17 | T1 |
| ES199 | ATCTGGGTGCTGCACACAAATCCG | | | | |
| ES200 | GTGATAGTGGAGCTTGCGGCAGTG | CNJ02860/CNJ02870 | 10 | 30-20 | T1 |
| ES66 | AATTCGAGAAGAGCGGGGTCATGAATACC | | | | |
| ES67 | AAATAAGGTGACAGAAAGCTGAGGAGGGC | CNG01990/CNG02000 | 7 | 37-01 | T1 |
| ES68 | TTACTTCCCGTTGCACTGTCATCTTCTCC | | | | |
| ES69 | ACTATAGGCTGACTGAGAGAGTAGAGCGC | CNH01775/CNH01780 | 8 | 37-01 | T1 |
| ES64 | CTGAATGTTAAGTGCAGCGTAATGAGCCG | | | | |
| ES65 | TCAATTCCAGTGGATAGCGTACGTTACCG | CNJ00540/CNJ00550 | 10 | 37-01 | T1 |
| ES70 | CTGGACAACCTCCACCTTTGTTTCTTTGC | | | | |
| ES71 | TCTGCTTCTTGTCCATCCATTTCTTCCCC | CNJ02660/CNJ02670 | 10 | 37-01 | T1 |
| ES116 | ATTAGAAGCTACCCTTGTTGTCGTCGTCG | | | | |
| ES117 | TGTGAGCAATTCGACATTGGGCATAAACG | CNA06700 | 1 | 37-01 | Tcn12 |
| ES114 | TAAACGAATTGTCACTAGGTCGTGGAGCC | | | | |
| ES115 | TGGCAATATTTGGTCCCGCTATCTGATCC | CNB03750 | 2 | 37-01 | Tcn12 |
| ES74 | ACTTACTACTGCACTCTCAATGGTGTGGC | | | | |
| ES75 | ACCGAATTTAGGGATGAGGATCTTGGTGC | CNA06050/CNA06060 | 1 | 37-02 | T1 |
| ES72 | CCTGCTTTTGCTTATGCCCAATCATACCC | | | | |
| ES73 | GTCTCTTCCCACTCTCCTTCTGTCTTTGC | CNA07360/CNA07370 | 1 | 37-02 | T1 |
| ES118 | TAGCCAACTGCGGATTGTTTGTTACAAGC | | | | |
| ES119 | ATTGCTGTTTCTCCGACTCATACAGTCCC | CNA07360/CNA07370 | 1 | 37-02 | T1 |
| ES76 | GACGTTGGCTGCCGTAATAATCTTTGTCC | | | | |
| ES77 | TCGATGGAATGGAAGGAAAGACACTGACG | CNB02690/CNB02680 | 2 | 37-02 | T1 |

| | | | | | |
|---|---|---|---|---|---|
| ES211 | GTCGCAGGTGCAGTTACCCACTTG | CNC05440 | 3 | 37-17 | Tcn12 |
| ES212 | TTCGATCACTTCCCTCCCATCCCC | | | | |
| ES213 | CAAGTCGAAGTGGGCTTGACCGTG | CNE03680 | 5 | 37-17 | Tcn12 |
| ES214 | AGGTGGTATCTGATGGGCGTCGTC | | | | |
| ES201 | ATGTGAGCCGCGTGATCCAAAAGG | CNB02900/CNB02910 | 2 | 37-18 | T1 |
| ES202 | CGGTTGAAACACGGCAACAAGCAC | | | | |
| ES150 | TTCTGAGCTCTGGGCATCCATCCC | CNN00910/CNN00920 | 14 | 37-18 | T1 |
| ES151 | GTCCGCAAGAGCATGAGCACTACG | | | | |
| ES215 | TTTGGTCTTGAGAGACGGCGGAGG | CNI04250 | 9 | 37-18 | Tcn12 |
| ES216 | GTACCCGAAAACCGCTCGATGTCG | | | | |
| ES217 | GCCGCCTCTCTTCATGTCGGTTTC | CNI02060 | 9 | 37-18 | Tcn12 |
| ES218 | ACCATGTGTGTACGGGATTGGTGC | | | | |
| ES154 | CGTACGTTCTCGTCGTGCTTGACC | CNA01360/CNA01370 | 1 | 37-19 | T1 |
| ES155 | TGGGATCAGGCGGATGAAGTCGAG | | | | |
| ES152 | CGCTTATGCAAGACCAGCCTCGAC | CNA08090/CNA08100 | 1 | 37-19 | T1 |
| ES153 | GCTGACTAGCGCGAGAAACAGGTG | | | | |
| ES156 | ATTGCGCAGAGGAGGTAGGAGACG | CNF01870/CNF01880 | 6 | 37-19 | T1 |
| ES157 | AGACGATGGTTCGCGCACATAAGC | | | | |
| ES158 | CGCTCAAAGGCTCGAAGTATCGCC | CNM02270/CNM02280 | 13 | 37-19 | T1 |
| ES159 | ACTGGTGGAGGTGATTCCGGTGTC | | | | |
| ES219 | AAACGAGCGCACGTACCAACATCG | CNF04770 | 6 | 37-19 | Tcn12 |
| ES220 | GGGTGGGGTATCACTTGCCATTGC | | | | |
| ES221 | CACAGTTGGTGTGGCCAGATCAGC | CNJ03330 | 10 | 37-19 | Tcn12 |
| ES222 | CGGCAAGCTGATCTCAGGAACGTC | | | | |
| ES160 | CGCTCAGTGCCGGTGTCAGTATTG | CNA03220/CNA03230 | 1 | 37-20 | T1 |
| ES161 | GCGTCTGGTCCAACGATATTGGCC | | | | |
| ES223 | AGAGAGCGTCGGAAGATTCGGGTG | CNA08120 | 1 | 37-20 | Tcn12 |
| ES224 | TGGAGCACAGGCAAATTGTCGAGG | | | | |
| ES225 | ACGCCAGCATTTATGACGACCACG | CNE05240 | 5 | 37-20 | Tcn12 |
| ES226 | AAGAACCTCATCCTGCGGCGATTG | | | | |
| ES164 | GAGAGCAAGCTCGTCAACGTCGTC | CND00300/CND00310 | 4 | 37-21 | T1 |
| ES165 | TTGCTTGGTGGCATTGTCTGACGG | | | | |
| ES166 | AGAAGCCTACATACCGGGCGTCAG | CND03920/CND03930 | 4 | 37-21 | T1 |
| ES167 | ACTCGTGCCCAGGTTCCCTAAAGG | | | | |
| ES168 | CGTGAAGCGCATGGGAACTGTAGG | CNI01880/CNI01890 | 9 | 37-21 | T1 |
| ES169 | GGAGCATCGAACCATCCGTCGTTG | | | | |
| ES227 | CAAAGGAGTGCCAGTCCGAGTTGC | CND04840 | 4 | 37-21 | Tcn12 |
| ES228 | CGAGTGTTGCGCAGGTCTACCATG | | | | |
| ES170 | ATCACTCACTCCTGGCCCAAGAGC | CNA00660/CNA00670 | 1 | 37-22 | T1 |
| ES171 | CTCAAAAGGAGGCGGGTTGACACG | | | | |
| ES172 | CTTCAACCCTTGCCACGTCTGCTC | CNB02545/CNB02550 | 2 | 37-22 | T1 |
| ES173 | TGCAGAAACAGTGACTGTGCGTCG | | | | |
| ES174 | GACTCGCAACCGATCCAGAAGTGC | CNC04810/CNC04820 | 3 | 37-22 | T1 |
| ES175 | AGAAGCGGAACCTGCGTGTGAAAC | | | | |
| ES176 | GCTTGTCCCAAGCTGCTCGTATGG | CNE03310/CNE03320 | 5 | 37-22 | T1 |
| ES177 | CCGTACCATTCTCGCACTCGCATG | | | | |

| ID | Full Name | 5'-3' DNA Sequence | Probe size (bp) | Source |
|---|---|---|---|---|
| ES130 | RetroF Cnl1 | AGCAGCACATCATCAAGCTC | 844 | Janbon G. et al., *Fungal Genet Biol.* 2010. |
| ES131 | RetroR Cnl1 | TAAATGACGCGGTTGATGGC | | |
| ES180 | Tcn12 Probe B F | TGAGAAAGCCATTCATTCGGCCGG | 537 | This study |
| ES181 | Tcn12 Probe B R | ATCCACTGGCGAGCCCTTCTACAC | | |
| JW83 | T1 Probe F | GTCGACGAGAGGAAATCTCATACTGTAC | 388 | This study |
| JW84 | T1 Probe R | CAGAGAGGACATCGGCAGCGG | | |

| ID | Target | 5'-3' DNA Sequence | Source |
|---|---|---|---|
| AG34 | GPD1 | GTCTCCACTGATTTCATTGGCTCTAC | JOHE44120; Fu et al. PLoS Genetics 2019 |
| AG35 | GPD1 | GTAACCATACTCATTGTCATACCAGCT | JOHE44121; Fu et al. PLoS Genetics 2019 |
| AG36 | GPD1 | GTAACCATACTCATTGTCATACTAAAAG | This study |
| AG37 | CNH01775 | CAGAAATTACAAGACGAACAGGACG | |
| AG38 | CNH01775 | GTGAAGCAACCGCTGGACTATC | |
| AG41 | CNH01780 | GCACTTATCCAAGGCCCTCC | |
| AG42 | CNH01780 | AGATCGGTACCCTCCGTTCAG | |
| AG45 | CNJ00540 | ATCAAGCCAGTCCCAGCATC | |
| AG46 | CNJ00540 | ATTGGCTTGGGTCTCAGCTC | |
| AG63 | CNJ00550 | GGCGTTCACCAAACCAACTG | |
| AG64 | CNJ00550 | TCCAAGAACGAATAGGGGCCA | |
| AG71 | CNC00200 | CTGGTGCAAAGATTCCCCCG | |
| AG72 | CNC00200 | CTCGAACCCCACACCTTCCTC | |
| AG75 | CNC00190 | ATGCGGCCACTTTTACCGAG | |
| AG76 | CNC00190 | GGTCGGCCATTGTTCTGAAGG | |
| AG81 | CNC05120 | TCAAGGCAAGCCAATGCTGG | |
| AG82 | CNC05120 | GCCGTCCGCTGAAGTTTGG | |
| AG87 | CNC05130 | GCTTCAATCCGCAAGCAAACT | |
| AG88 | CNC05130 | TTCCAGTCGATCTCGAAACGC | |
| AG91 | CNK01950 | AGGCCTTGGACTTACAGGAAAAC | |
| AG92 | CNK01950 | CTGGGAGTTGCCTGACAAGG | |
| AG97 | CNK01960 | AGGAGGAATGCCCATGGTTG | |
| AG98 | CNK01960 | AACATCCGCAACGGTCTTGG | |
| AG105 | CNM00880 | CGAGGACGAACTTCAGGACG | |
| AG106 | CNM00880 | TGGCTCGCTTATTGGCCTTG | |
| AG109 | CNM00890 | ACAGGACCAATGTCGCACAA | |
| AG110 | CNM00890 | TGATCGGCCCACACATTCC | |
| AG143 | 5'-end Cnl1 | GGCTCTTCCCTAGTCGCTTG | |
| AG144 | 5'-end Cnl1 | CAGTATTGGAGGGAGGGCAG | |

**Table S5.** Sequence variations detected in Illumina-sequenced TA lines.

| TA line | Type | Chromosome | Position | Reference Allele | Alternative Allele | Location | Amino Acid Change |
|---|---|---|---|---|---|---|---|
| 30-01 | SNP | XL280_Chr05 | 15362 | G | C | Start of CNE05200 | |
| 30-01 | SNP | XL280_Chr09 | 239663 | G | A | Third exon of CNI00980 | synonymous |
| 30-01 | SNP | XL280_Chr12 | 1064289 | T | C | intergenic, upstream CNL06555 | |
| 30-01 | SNP | XL280_Chr03 | 1823885 | A | G | intergenic | |
| 30-01 | SNP | XL280_Chr14 | 288677 | G | A | Third exon of CNN00890 | R → Q |
| 30-02 | SNP | XL280_Chr01 | 10750 | G | A | Last exon of CNA00030 | G → S |
| 30-02 | SNP | XL280_Chr01 | 219916 | C | G | Last exon of CNA00760 | P → A |
| 30-02 | SNP | XL280_Chr04 | 1685043 | C | A | Exon of CND06090 | S → N |
| 30-02 | SNP | XL280_Chr08 | 497081 | A | T | Start of CNH01460 and CNH01480 | |
| 30-02 | SNP | XL280_Chr10 | 566700 | C | G | First exon on CNJ01930 | G → D |
| 30-02 | SNP | XL280_Chr13 | 722802 | T | A | intergenic | |
| 30-02 | ins | XL280_Chr02 | 930641 | AATA | AATATA | intergenic | |
| 30-02 | ins | XL280_Chr06 | 57476 | CTATTA | CTATTATTA | Downstream of CNC02090 | |
| 30-03 | SNP | XL280_Chr03 | 1712429 | C | T | intergenic, upstream of CNC05750 | |
| 30-03 | SNP | XL280_Chr11 | 862569 | T | C | intronic, between the third and fourth exons of CNK02850 | |
| 30-03 | del | XL280_Chr02 | 930643 | TAC | T | intergenic | |
| 30-16 | SNP | XL280_Chr02 | 1112 | C | T | End of First exon in CNB00010 | H → Y |
| 30-16 | SNP | XL280_Chr04 | 556359 | C | T | Sixth exon of CND02040 | Q → STOP |
| 30-16 | SNP | XL280_Chr05 | 1151182 | C | T | Within CNE04070 (hypothetical protein) | |
| 30-16 | SNP | XL280_Chr08 | 168992 | C | T | intronic in CNH02510, between exons four and five | |
| 30-16 | SNP | XL280_Chr08 | 573639 | G | A | First exon of CNH01250 | G → S |
| 30-16 | SNP | XL280_Chr11 | 191418 | T | C | intergenic | |
| 30-16 | SNP | XL280_Chr12 | 422229 | G | A | Middle of seventh exon of gene CNL04260 | synonymous |
| 30-16 | ins | XL280_Chr02 | 976982 | CATATATATAT | CATATATATATAT | 5' UTR CNB03180 | |
| 30-16 | ins | XL280_Chr08 | 442476 | ATT | ATTTT | intergenic | |
| 30-16 | del | XL280_Chr11 | 39688 | GAAAAAAAAAAA | GAAAAAAAAAA | 3' UTR CNK00120 and upstream of CNK00110 | |
| 30-17 | SNP | XL280_Chr01 | 92717 | G | A | 5' UTR CNA00290 | |
| 30-17 | SNP | XL280_Chr01 | 802397 | C | G | Fourth exon of CNA03070 | synonymous |
| 30-17 | SNP | XL280_Chr05 | 170156 | G | A | Start of third exon of CNE00630 | A → V |
| 30-17 | SNP | XL280_Chr09 | 878884 | A | C | intergenic | |
| 30-17 | SNP | XL280_Chr09 | 1148096 | T | G | Middle of CNI04310 | |
| 30-17 | SNP | XL280_Chr10 | 489241 | T | C | intergenic, between CNJ01650 and CNJ01660 | |
| 30-17 | ins | XL280_Chr12 | 959798 | ATT | ATTT | intergenic | |
| 30-18 | SNP | XL280_Chr01 | 2295566 | G | A | Fifth exon of CNA08320 | A → V |
| 30-18 | SNP | XL280_Chr04 | 1184333 | C | T | intergenic | |
| 30-18 | SNP | XL280_Chr07 | 520652 | G | C | 5' UTR CNG01820 | |
| 30-18 | SNP | XL280_Chr08 | 643616 | G | A | intergenic | |
| 30-18 | SNP | XL280_Chr11 | 8490 | T | A | intergenic, upstreamof CNK00020 | |
| 30-18 | SNP | XL280_Chr14 | 77727 | C | T | 3' UTR CNN00175 | |
| 30-18 | ins | XL280_Chr01 | 535657 | CA | CAA | intergenic | |
| 30-18 | del | XL280_Chr12 | 1060876 | AG | A | 3' UTR CNL06555 | |
| 30-18 | ins | XL280_Chr02 | 930640 | TAA | TAAA | intergenic | |
| 30-19 | SNP | XL280_Chr03 | 211100 | C | T | 5' UTR CNC00705 | |
| 30-19 | SNP | XL280_Chr03 | 1353827 | A | T | 5' UTR CNC04440 | |
| 30-19 | SNP | XL280_Chr05 | 529530 | G | A | Intronic, between the 12th and 13th exons of CNE01970 | |
| 30-19 | SNP | XL280_Chr05 | 529532 | T | A | Intronic, between the 12th and 13th exons of CNE01970 | |
| 30-19 | SNP | XL280_Chr05 | 529533 | A | T | Intronic, between the 12th and 13th exons of CNE01970 | |
| 30-19 | SNP | XL280_Chr06 | 270095 | C | G | 3' UTR CNF00810 | |
| 30-19 | SNP | XL280_Chr09 | 993670 | A | G | First exon of CNI03680 | F → S |
| 30-19 | SNP | XL280_Chr12 | 672917 | A | G | intergenic, downstream of CNL05120 | |
| 30-19 | del | XL280_Chr01 | 1701439 | AATATAT | AATAT | 3' UTR CNA06220 | |
| 30-19 | ins | XL280_Chr12 | 984440 | TAA | TAAA | 5' UTR CNL06320 | |
| 30-20 | SNP | XL280_Chr01 | 296665 | G | T | Third exon of CNA01100 | A → D |
| 30-20 | SNP | XL280_Chr01 | 1894226 | C | A | intergenic, downstream of CNA06940 | |
| 30-20 | SNP | XL280_Chr01 | 1894227 | A | T | intergenic, downstream of CNA06940 | |
| 30-20 | SNP | XL280_Chr02 | 773886 | T | C | Intronic, between the 14th and 15th exons of CNB02580 | |
| 30-20 | SNP | XL280_Chr03 | 361184 | T | G | Intronic, between the eighth and ninth exons of CNC01260 | |
| 30-20 | SNP | XL280_Chr03 | 818514 | A | T | intergenic | |
| 30-20 | SNP | XL280_Chr04 | 69985 | C | T | intergenic, upstream of CND00210 | |
| 30-20 | SNP | XL280_Chr05 | 1321956 | T | A | 3' UTR CNE04710 | |
| 30-20 | SNP | XL280_Chr07 | 1266968 | G | A | Second exon of CNG04300 | D → N |
| 30-20 | SNP | XL280_Chr08 | 461733 | C | T | intergenic | |
| 30-20 | SNP | XL280_Chr14 | 45058 | A | G | First exon of CNN00120 | synonymous |
| 30-20 | del | XL280_Chr12 | 959796 | AC | A | intergenic | |
| 30-20 | SNP | XL280_Chr01 | 1012449 | G | C | 3' UTR CNA03770 or 2nd to last exon of CNA03780 | R→ G |
| 30-20 | ins | XL280_Chr07 | 675124 | TAA | TAAA | intergenic | |

**Table S5 cont.** Sequence variations detected in Illumina-sequenced TA lines.

| TA line | Type | Chromosome | Position | Reference Allele | Alternative Allele | Location | Amino Acid Change |
|---|---|---|---|---|---|---|---|
| 37-01 | SNP | XL280_Chr03 | 1153796 | C | T | Start of second exon of CNC03640 | synonymous |
| 37-01 | SNP | XL280_Chr03 | 1258899 | C | A | Start of the first exon of CNC04030 | A → S |
| 37-01 | SNP | XL280_Chr05 | 750241 | C | T | First exon of CNE02840 | H → Y |
| 37-01 | SNP | XL280_Chr08 | 812409 | T | A | 5' UTR CNH00440 | |
| 37-01 | SNP | XL280_Chr08 | 812416 | T | A | 5' UTR CNH00440 | |
| 37-01 | SNP | XL280_Chr11 | 31047 | T | C | First exon of CNK00090 | synonymous |
| 37-01 | SNP | XL280_Chr11 | 192953 | C | T | intergenic | |
| 37-01 | SNP | XL280_Chr13 | 588580 | G | C | Intronic, between the second and third exons of CNM01790 | |
| 37-01 | ins | XL280_Chr06 | 550147 | GGTCG | GGTCGTCG | intergenic | |
| 37-01 | del | XL280_Chr09 | 83065 | TAAA | TAA | 5'UTR CNI00350 | |
| 37-02 | SNP | XL280_Chr01 | 2187372 | C | T | End of second exon of CNA07900 | synonymous |
| 37-02 | SNP | XL280_Chr02 | 146365 | G | C | intergenic, downstream of CNB00500 | |
| 37-02 | SNP | XL280_Chr03 | 984134 | C | G | intergenic | |
| 37-02 | SNP | XL280_Chr05 | 884767 | A | T | intergenic | |
| 37-02 | SNP | XL280_Chr12 | 547270 | A | G | 3' UTR CNL04680 | |
| 37-02 | ins | XL280_Chr02 | 56981 | CAAAAAA | CAAAAAAA | Right side of unkown gene CNB00180 | |
| 37-02 | ins | XL280_Chr04 | 740203 | ACAG | ACAGCAG | 3' UTR CND02690 | |
| 37-02 | ins | XL280_Chr07 | 984812 | ATAG | ATAGTAG | intergenic | |
| 37-03 | SNP | XL280_Chr01 | 1258883 | C | T | End of second exon of CNA04760 | A → S |
| 37-03 | SNP | XL280_Chr02 | 1516464 | C | G | second exon of CNB05310 | synonymous |
| 37-03 | SNP | XL280_Chr03 | 1429233 | C | T | 5' UTR CNC04720 | |
| 37-03 | SNP | XL280_Chr04 | 1209022 | G | T | intronic, between the second and third exons of CND04340 | |
| 37-03 | SNP | XL280_Chr07 | 59136 | G | A | intergenic | |
| 37-03 | SNP | XL280_Chr12 | 414271 | C | T | End of first exon of CNL04230 | E → K |
| 37-03 | SNP | XL280_Chr13 | 814001 | C | G | Middle of second exon of CNM02550 | synonymous |
| 37-16 | SNP | XL280_Chr01 | 372508 | C | G | Last exon of CNA01360 | A → G |
| 37-16 | SNP | XL280_Chr01 | 1995534 | T | C | 3' UTR CNA07270 | |
| 37-16 | SNP | XL280_Chr01 | 2105474 | C | T | Start of seventh exon of CNA07650 | synonymous |
| 37-16 | SNP | XL280_Chr01 | 2135960 | A | C | intergenic | |
| 37-16 | SNP | XL280_Chr03 | 1918108 | T | G | Sixth exon of CNC06530 | V → G |
| 37-16 | SNP | XL280_Chr03 | 1918109 | T | G | Sixth exon of CNC06530 | V → G |
| 37-16 | SNP | XL280_Chr03 | 1918110 | C | G | Sixth exon of CNC06530 | R → G |
| 37-16 | SNP | XL280_Chr04 | 924944 | C | T | Last exon of CND03390 | A → V |
| 37-16 | SNP | XL280_Chr06 | 1245185 | T | C | Third exon of CNF04340 | synonymous |
| 37-16 | SNP | XL280_Chr07 | 499876 | C | T | Start of last exon of CNG01730 | L → F |
| 37-16 | SNP | XL280_Chr07 | 627715 | G | A | intergenic | |
| 37-16 | SNP | XL280_Chr07 | 820480 | C | T | 5' UTR CNG02950 | |
| 37-16 | SNP | XL280_Chr09 | 554445 | G | A | Third exon of CNI02020 | S → N |
| 37-16 | SNP | XL280_Chr11 | 717431 | A | T | intergenic | |
| 37-16 | SNP | XL280_Chr12 | 61390 | T | G | Start of CNH03240 | |
| 37-16 | SNP | XL280_Chr12 | 432012 | C | T | Fifteenth exon of CNL04300 | S → L |
| 37-16 | SNP | XL280_Chr13 | 293825 | A | G | End of third exon of CNM00850 | synonymous |
| 37-16 | del | XL280_Chr01 | 1741511 | GCGCTCGAAT | G | Three amino acid deletion in first exon of CNA06390 | Non-synonymous |
| 37-17 | SNP | XL280_Chr02 | 1554542 | C | T | End of the fourth exon of CNB05450 | T → I |
| 37-17 | SNP | XL280_Chr04 | 1429507 | C | T | intronic, between the third and fourth exons of CND05210 | |
| 37-17 | SNP | XL280_Chr09 | 249373 | A | C | intronic, betweeen the second and thrid exon of CNI01010 | |
| 37-17 | SNP | XL280_Chr10 | 192884 | T | A | 5' UTR CNJ00650 | |
| 37-17 | del | XL280_Chr02 | 818765 | CA | C | intergenic | |
| 37-18 | SNP | XL280_Chr03 | 655336 | C | T | Twelfth exon of CNC02260 | synonymous |
| 37-18 | SNP | XL280_Chr04 | 1532615 | C | T | 5' UTR CND05660 | |
| 37-18 | SNP | XL280_Chr06 | 1387880 | G | A | Intronic, between the third and fourth exons of CNF04830 | |
| 37-18 | SNP | XL280_Chr09 | 190234 | A | C | Second exon of CNI00790 | I → L |
| 37-18 | ins | XL280_Chr02 | 930637 | AAATAATA | AAATAATAATA | intergenic | |
| 37-19 | SNP | XL280_Chr02 | 1025015 | G | A | First exon of CNB03350 | R → C |
| 37-19 | SNP | XL280_Chr04 | 891040 | G | C | Middle of the last exon of CND03290 | E → D |
| 37-19 | SNP | XL280_Chr05 | 1468894 | G | A | intergenic | |
| 37-19 | SNP | XL280_Chr12 | 243336 | T | G | Intergenic, downstream of CNL03840 | |
| 37-19 | SNP | XL280_Chr13 | 105634 | C | T | Middle of last exon of CNM00300 | synonymous |
| 37-19 | del | XL280_Chr05 | 1465624 | TTAGGGG | T | intergenic | |
| 37-19 | del | XL280_Chr10 | 1095668 | TTAGGGG | T | intergenic | |
| 37-19 | ins | XL280_Chr11 | 3415 | AAC | AACAC | intergenic | |
| 37-19 | ins | XL280_Chr12 | 984442 | AT | ATT | 5' UTR CNL06320 | |
| 37-19 | SNP | XL280_Chr08 | 942065 | C | A | intergenic, near telomeric end of chromosome | |
| 37-19 | SNP | XL280_Chr08 | 942141 | T | C | intergenic, near telomeric end of chromosome | |
| 37-20 | SNP | XL280_Chr04 | 1500543 | G | T | Middle of first exon of CND05510 | P → Q |
| 37-20 | SNP | XL280_Chr05 | 928064 | C | T | Third exon of CNE03250 | P → S |
| 37-20 | SNP | XL280_Chr06 | 402347 | T | G | Start of the first exon of CNF01360 | Q → H |
| 37-20 | SNP | XL280_Chr08 | 330959 | C | T | Middle of the last exon of CNH01920 | G → W |
| 37-20 | SNP | XL280_Chr09 | 176732 | T | C | Fourth exon of CNI00720 | D → G |
| 37-20 | del | XL280_Chr02 | 976993 | TA | T | 5' UTR CNB03180 | |
| 37-21 | SNP | XL280_Chr01 | 613931 | G | A | Eighth exon of CNA02350 | E → K |
| 37-21 | SNP | XL280_Chr11 | 194687 | T | A | intergenic | |
| 37-21 | del | XL280_Chr03 | 464160 | TAA | TA | intergenic | |
| 37-22 | SNP | XL280_Chr07 | 122589 | G | A | Last exon of CNG00470 | synonymous |
| 37-22 | SNP | XL280_Chr09 | 899517 | T | A | intergenic, upstream of CNI03320 | |
| 37-22 | SNP | XL280_Chr12 | 1046929 | C | T | 5' UTR CNL06500 | |
